# The relationship between EEG and fMRI connectomes is reproducible across simultaneous EEG-fMRI studies from 1.5T to 7T

**DOI:** 10.1101/2020.06.16.154625

**Authors:** Jonathan Wirsich, João Jorge, Giannarita Iannotti, Elhum A Shamshiri, Frédéric Grouiller, Rodolfo Abreu, François Lazeyras, Anne-Lise Giraud, Rolf Gruetter, Sepideh Sadaghiani, Serge Vulliémoz

**Affiliations:** EEG and Epilepsy Unit, University Hospitals and Faculty of Medicine of Geneva, Geneva, Switzerland; Laboratory for Functional and Metabolic Imaging, École Polytechnique Fédérale de Lausanne, Lausanne, Switzerland; Systems Division, Swiss Center for Electronics and Microtechnology (CSEM), Neuchâtel, Switzerland; Swiss Center for Affective Sciences, University of Geneva, Geneva, Switzerland; ISR-Lisboa/LARSyS and Department of Bioengineering, Instituto Superior Técnico – Universidade de Lisboa, Lisbon, Portugal; Coimbra Institute for Biomedical Imaging and Translational Research (CIBIT), Institute for Nuclear Sciences Applied to Health (ICNAS), University of Coimbra, Coimbra, Portugal; Department of Radiology and Medical Informatics, University of Geneva, Geneva, Switzerland; Department of Neuroscience, University of Geneva, Geneva, Switzerland; Department of Radiology, University of Lausanne, Lausanne, Switzerland; Beckman Institute, University of Illinois at Urbana-Champaign, Urbana, IL, USA; Psychology Department, University of Illinois at Urbana-Champaign, Urbana, IL, USA

## Abstract

Both electroencephalography (EEG) and functional Magnetic Resonance Imaging (fMRI) are non-invasive methods that show complementary aspects of human brain activity. Despite measuring different proxies of brain activity, both the measured blood-oxygenation (fMRI) and neurophysiological recordings (EEG) are indirectly coupled. The electrophysiological and BOLD signal can map the underlying functional connectivity structure at the whole brain scale at different timescales. Previous work demonstrated a moderate but significant correlation between resting-state functional connectivity of both modalities, however there is a wide range of technical setups to measure simultaneous EEG-fMRI and the reliability of those measures between different setups remains unknown. This is true notably with respect to different magnetic field strengths (low and high field) and different spatial sampling of EEG (medium to high-density electrode coverage).

Here, we investigated the reproducibility of the bimodal EEG-fMRI functional connectome in the most comprehensive resting-state simultaneous EEG-fMRI dataset compiled to date including a total of 72 subjects from four different imaging centers. Data was acquired from 1.5T, 3T and 7T scanners with simultaneously recorded EEG using 64 or 256 electrodes. We demonstrate that the whole-brain monomodal connectivity reproducibly correlates across different datasets and that a moderate crossmodal correlation between EEG and fMRI connectivity of r≈0.3 can be reproducibly extracted in low- and high-field scanners. The crossmodal correlation was strongest in the EEG-β frequency band but exists across all frequency bands. Both homotopic and within intrinsic connectivity network (ICN) connections contributed the most to the crossmodal relationship.

This study confirms, using a considerably diverse range of recording setups, that simultaneous EEG-fMRI offers a consistent estimate of multimodal functional connectomes in healthy subjects that are dominantly linked through a functional core of ICNs across spanning across the different timescales measured by EEG and fMRI. This opens new avenues for estimating the dynamics of brain function and provides a better understanding of interactions between EEG and fMRI measures. This observed level of reproducibility also defines a baseline for the study of alterations of this coupling in pathological conditions and their role as potential clinical markers.

## Introduction

The brain is a complex system of interacting neurons continuously communicating with each other. This intrinsic brain functional connectivity (FC) has been shown to be organized in macro-scale patterns of interconnected regions, the so-called intrinsic connectivity networks (ICNs) (Biswal et al., 1995; Fox et al., 2005; Greicius et al., 2003). Though originally described for fMRI (FC_fMRI_) data, those ICNs have also been found using Magnetoencephalography (MEG, FC_MEG_) (Brookes et al., 2011) and Electroencephalography (EEG, FC_EEG_) (Abreu et al., 2020; de Pasquale et al., 2010; de Pasquale et al., 2012; Finger et al., 2016; Wirsich et al., 2017). Expanding this initially network-specific view, cross-modal studies showed that FC in both electrophysiological (M/EEG) and BOLD signals are related when using static measures (averaged over periods >5 minutes) across the whole brain scale of resting-state recordings (Deligianni et al., 2014; Hipp and Siegel, 2015; Wirsich et al., 2017). Understanding the functional links between EEG- and fMRI-derived FC measures is a key question in current neuroimaging research, as it could help clarifying the neuronal substrates of both modalities, and of resting-state activity itself (Sadaghiani and Wirsich, 2020).

The relation between FC_fMRI_ and FC_M/EEG_, has also been shown to be mediated by the underlying white matter structural connectivity derived from diffusion MRI (SC_dMRI_) (Chu et al., 2015; Deligianni et al., 2019, 2016; Honey et al., 2009; Meier et al., 2016; Wirsich et al., 2017). Beyond the static relationship of EEG and fMRI to structure, we also observed crossmodal linked dynamics (1 minute sliding window) (Wirsich et al., 2020b). While most work focused on crossmodal agreement of FC, we have equally shown that the crossmodal connectivity can be split up into a common and complimentary connectivity profile across different timescales (Wirsich et al., 2020a). A crucial open question is whether crossmodal dissimilarities arise mainly from differences in data quality and acquisition setup, or represent differences arising from measuring complementary multimodal aspects of brain functional connectivity (Sadaghiani and Wirsich, 2020). In order to address this issue, data from independent research sites are needed. This would enable us to characterize the baseline relationship between connectivity derived from both modalities.

From a monomodal point of view, the test-retest reliability of FC_fMRI_ measures is well characterized (Noble et al., 2019) with a low intra class correlation (ICC=0.29) for individual connections and reproducible ICN estimations across sites (Badhwar et al., 2020). While some of the variance stems from different scanner systems (Han et al., 2006) or inter-individual differences (Amico and Goñi, 2018; Finn et al., 2015), the chosen post-processing of data also introduces variability of the results (Botvinik-Nezer et al., 2020; Carp, 2012). Due to this connectivity variability, it is important to estimate the reproducibility of this measure, as a baseline for subsequent measures of alterations in different normal and pathological conditions (De Vico Fallani et al., 2014).

The reproducibility of FC_M/EEG_ measures has been analyzed from different angles. Different measures of FC (Colclough et al., 2016) have shown highly correlated (topographically similar) connectomes while Coquelet et al. (2020) have shown that the crossmodal relationship of MEG and EEG connectivity is reproducible across different EEG forward models. Marquetand et al. (2019) observed good intersession test-retest reliability (ICC>0.67) for both modalities. From a frequency point of view, alpha has been shown to be the most reliable estimate across subjects (Colclough et al., 2016; Marquetand et al., 2019). While the majority of FC_fMRI_ work derives connectivity from correlating regional timecourses of the BOLD signal, no consensus has been reached yet for FC_M/EEG_, especially whether to use phase or amplitude coupling (Colclough et al., 2016; Sadaghiani and Wirsich, 2020). Both phase coupling using the imaginary part of the coherency (iCoh) (Nolte et al., 2004; Wirsich et al., 2017) and amplitude coupling using amplitude envelope correlation (AEC) (Brookes et al., 2011; de Pasquale et al., 2010; Deligianni et al., 2014; Hipp and Siegel, 2015) have been linked to FC_fMRI_. It has been argued that phase coupling might be closer to the underlying SC_dMRI_ and amplitude coupling is proposed to be more related to FC_fMRI_ (Engel et al., 2013). Nevertheless, most of the literature suggests a rather similar connectivity pattern of intrinsic brain activity on the whole brain scale when averaging over resting-state recordings lasting several minutes (Colclough et al., 2016; Mostame and Sadaghiani, 2020; Sadaghiani and Wirsich, 2020). Siems and Siegel (2020) observed highly correlated, but not identical connectivity between both types of measures, and the neurobiological interpretation of those differences have yet to be explored. Recently those changes have been linked to task-specific connectivity (Mostame and Sadaghiani, 2020), but ultimately the relevance of different connectivity measures for brain behavior and pathology is still an open question.

Electrophysiological measures suffer from an ill-posed problem, when reconstructing sparsely sampled sensor signals on the scalp into the three-dimensional brain space, resulting in source-activity leaking into different regions which can distort connectivity measures (Palva et al., 2018). The selection of an optimal brain parcellation can reduce the variability arising from regional crosstalk due to source leakage of the inverse solution (Farahibozorg et al., 2018). Given those limitations of electrophysiological measures, less work has been done to extract electrophysiological connectomes (Sadaghiani and Wirsich, 2020). However, no study has compared if the EEG or the fMRI connectome can be measured more reliably across subjects, especially in a simultaneous recording.

In the case of simultaneous EEG-fMRI, signal quality is diminished by 1) artifacts on fMRI data induced by the EEG electrodes interacting with the static magnetic field and the MR-pulses (Mullinger et al., 2008b) 2)gradient- and pulse-related EEG artifacts induced by the magnetic gradients on the EEG electrodes and cables (Abreu et al., 2018; Allen et al., 2000). At 7T, EEG-fMRI can be affected by stronger recording artifacts (Abreu et al., 2016; Jorge et al., 2015b; Mullinger et al., 2008a; Neuner et al., 2013). Overcoming those limitations would encourage analysis of the dynamic electrophysiological correlates of BOLD signal at much higher temporal and spatial resolutions (Meyer et al., 2019; Scheeringa and Fries, 2017). Though it is well documented that the data quality of EEG and fMRI depends on scanner field strength (Debener et al., 2008; Mullinger et al., 2008b), no data exists assessing the impact on the data reliability at the level of functional connectivity. The possibility to compensate for the larger artifacts at higher field (Jorge et al., 2015a) might be sufficient to achieve reliable EEG-fMRI connectivity measures.

To close the gap of unknown reproducibility of EEG-fMRI connectomes across experimental setups and to evaluate the suitability of a 7T EEG-fMRI setup for multimodal connectomics in this study we aim to:

1. compare the monomodal topographical similarity of FC_fMRI_ to FC_EEG_ derived from simultaneous EEG-fMRI across different imaging centers
2. characterize the reproducibility of crossmodal FC_fMRI_-FC_EEG_ relationship across heterogeneous setups and, for the first time, at 7T
3. characterize the robustness of the crossmodal relationship to methodological choices regarding the chosen brain parcellation and EEG connectivity measure
4. characterize the stability of the crossmodal relationship with respect to acquisition duration and number of subjects used for group averages
5. characterize the spatial contributions of individual connections to this crossmodal relationship and the topographical similarity of these contribution across different datasets

We will consider that the crossmodal relationship is reproducible if the monomodal measures are correlated across datasets and if the crossmodal relationship remains significant across all datasets. We further expect that the crossmodal relationship is robust to methodological choices. For example, we expect a robust crossmodal relationship to be significant for a range of methodological choices while the magnitude of the relationship might change. Our previous work suggest this correlation to be moderate around r~0.3 (Wirsich et al., 2020a, 2017). The reproducibility of monomodal measures, the crossmodal relationship and the robustness of the latter to methodological choices would strongly support the generalizability of concurrently recorded EEG-fMRI connectomes.

To do so, we combined simultaneous EEG-fMRI resting state acquisitions from 4 different centers totaling 72 subjects, and comprising recordings using a 1.5T, 3T and 7T MR scanner in combination with a 64- or 256-electrode EEG system.

## Methods

### Subjects and acquisition setup

We analyzed a total of 72 subjects divided up into 4 datasets: 16 subjects using a 64-channel EEG setup in a 1.5T MR-scanner (64Ch-1.5T), 26 subjects using a 64-channel EEG setup in a 3T MR-scanner (64Ch-3T), 21 subjects using a 256-channel EEG setup in a 3T MR-scanner (256Ch-3T) and 9 subjects using a 64 channel EEG setup in a 7T MR-scanner (64Ch-7T). Main differences between study paradigm, hardware and software setup are summarized in SI Table 1.

### Data set 1 (64Ch-1.5T)

16 subjects (6 females, mean age: 32.87, range 22-53) with no history of neurological or psychiatric illness were recorded. Ethical approval was given by local Research Ethics Committee (UCL Research ethics committee, project ID: 4290/001) and informed consent was obtained from all subjects (Deligianni et al., 2016, 2014). In each subject one run of 10 min 48 seconds resting-state simultaneous EEG-fMRI was acquired. Subjects were asked not to move, to remain awake and fixate on a white cross presented on a black background. MRI was acquired using a 1.5 Tesla MR-scanner (Siemens Avanto). The fMRI scan comprised of the following parameters: GRE-EPI sequence, TR=2160, TE=30ms, 30 slices, 210×210mm Field of View, voxel size 3.3×3.3×4.0mm^3^ (1mm gap), flip angle 75°, total of 300 volumes. The subjects’ head was immobilized using a vacuum cushion during scanning. Additionally, an anatomical T1-weighted image was acquired (176 sagittal slices, 1.0×1.0×1.0 mm, TA=11min). EEG was acquired using two 32-channel MR-compatible amplifiers (BrainAmp MR, sampling rate 1kHz), 63 electrodes (BrainCap MR, Gilching, Germany), referenced to FCz, 1 ECG electrode. The scanner clock was time-locked with the amplifier clock (Mandelkow et al., 2006). The MR-compatible amplifier was positioned behind outside the bore behind the head of the subject.

### Data set 2 (64Ch-3T)

26 healthy subjects (8 females, mean age 24.39, age range 18-31) with no history of neurological or psychiatric illness were recorded. Ethical approval was given by local Research Ethics Committee (CPP Ile de France III) and informed consent was obtained from all subjects (Sadaghiani et al., 2010). In each subject, 3 runs of 10 minutes (total 30 mins) resting-state simultaneous EEG-fMRI were acquired. Subjects were asked not to move and to remain awake and keep their eyes closed during the resting-state scan. For three subjects, one out of the three rest sessions was excluded due to insufficient EEG quality. The resting-state sessions were part of a study with two additional naturalistic film stimuli of 10 minutes not analyzed in the current study, and acquired after resting runs 1 and 2 of the resting state as described in Morillon et al. (Morillon et al., 2010). MRI was acquired using a 3 Tesla MR-scanner (Siemens Tim-Trio). The fMRI scan comprised of the following parameters: GRE-EPI sequence, TR=2000ms, TE=50ms, 40 slices, 192×192mm Field of View, voxel size 3×3×3mm^3^, flip angle 78°, total of 150 volumes (total all sessions 450 volumes). An anatomical T1-weighted image was acquired (176 sagittal slices, 1.0×1.0×1.0 mm, TA=7min). EEG was acquired using two 32-channel MR-compatible amplifiers (BrainAmp MR, sampling rate 5kHz), 62 electrodes (Easycap, Herrsching, Germany), referenced to FCz, 1 ECG electrode, and 1 EOG electrode. The scanner clock was time-locked with the amplifier clock (Mandelkow et al., 2006). The MR-compatible amplifier was positioned behind outside the bore behind the head of the subject.

### Data set 3 (256Ch-3T)

21 healthy subjects (7 females, 32.13, age range 24-47) with no history of neurological or psychiatric illness were recorded. Ethical approval was given by local Research Ethics Committee (Ethics committee of Geneva) and informed consent was obtained from all subjects. A subgroup of this cohort has been already analyzed by (Iannotti et al., 2015). In each subject one run of 4min 58.5s resting-state simultaneous EEG-fMRI were acquired. Five subjects had a longer recording of 19min 52s and one subject had a recording of 9min 56s, in all those cases the total run was analyzed. Subjects were asked not to move and to remain awake and keep their eyes closed during the resting-sate scan. MRI was acquired using a 3 Tesla MR-scanner (Siemens Magnetom Trio). The fMRI scan comprised the following parameters: GRE-EPI sequence, TR=1990ms, TE=30ms, 32 slices, 192×192mm Field of View, voxel size 3×3×3.75mm^3^, flip angle 90°, total of 150 volumes. Additionally, an anatomical T1-weighted image was acquired (176 sagittal slices, 1.0×1.0×1.0 mm, TA=7min). EEG was acquired using a 258-channel MR-compatible amplifier (Electrical Geodesic Inc., Eugene, OR, USA, sampling rate 1kHz), 256 electrodes (Geodesic Sensor Net 256), referenced to Cz, 2 ECG electrodes. The scanner clock was time-locked with the amplifier clock (Mandelkow et al., 2006). An elastic bandage was pulled over the subjects’ head and EEG cap to assure the contact of electrodes on the scalp. The MR-compatible amplifier was positioned to the left of the subject and EEG and ECG cables were passed through the front end of the bore.

### Data set 4 (64Ch-7T)

9 healthy subjects (4 females, mean age 23.56, age range 22-26) with no history of neurological or psychiatric illness were recorded. Ethical approval was given by the local Research Ethics Committee (CER-VD) and informed consent was obtained from all subjects (Jorge et al., 2019). In each subject 1 run of 8 minutes resting-state simultaneous EEG-fMRI was acquired. Subjects were asked not to move in the MR scanner and to keep their eyes open during the resting-state scan, fixating on a small red cross presented on a grey background, to minimize head and eye movements. Padding was also used to further restrict motion. MRI was acquired using a 7 Tesla head MR-scanner (Siemens Magnetom). The fMRI scan was performed using a simultaneous multi-slice (SMS) GRE-EPI sequence (3× SMS and 2× in-plane GRAPPA accelerations), with TR=1000ms, TE=25ms, 69 slices, 220×220mm Field of View, voxel size 2.2×2.2×2.2 mm, flip angle 54°, and a total of 480 volumes. A short EPI acquisition (5 volumes) with reversed phase encoding direction was also performed, for image distortion correction. Additionally, an anatomical T1-weighted image was acquired (160 sagittal slices, 1.0×1.0×1.0mm, TA = 10min). EEG was acquired using two 32-channel MR-compatible amplifiers (BrainAmp MR, sampling rate 5kHz), and a 63-electrode (EasyCap, Herrsching, Germany), referenced to FCz, 1 ECG electrode, connected via 12-cm bundled cables to reduce artifact contributions (Jorge et al., 2015b). Four of the 64 electrodes (T7, T8, F5 and F6) were customized to serve as motion artifact sensors (Jorge et al., 2015a). A total of 59 electrodes therefore remained dedicated to EEG recording. The scanner clock was time-synchronized with the amplifier clock (Mandelkow et al., 2006)). The MR-compatible amplifier was positioned inside the bore behind the head of the subject.

### Analysis

As data acquisition already has a considerable number of varying parameters that might affect the final FC estimation, we stress here that also for the analysis we did not strictly control every processing step. The rationale of this was that independent of setup and postprocessing, the different datasets should generalize across a family of analyses (Botvinik-Nezer et al., 2020) and result in reproducible monomodal and crossmodal measures. Applied to this study, the crossmodal relationship can be considered reproducible if it is robust to different hardware setups and preprocessing approaches. The different starting points across datasets given by the different EEG and fMRI equipment described above make it hard to acquire perfectly unbiased recordings across datasets. In consequence, our approach here was to optimize the analysis of each dataset to obtain the best possible signal quality (E.g. varying with field strength, several fMRI parameters are affected: e.g. optimal TE and thereby the time available to execute the EPI readout; techniques like GRAPPA and SMS-EPI - like used in the 7T data - become more or less effective, the spatial resolution vs. physiological noise relationship changes etc.). Regarding EEG side, as pointed out later, the channel geometry (Iannotti et al., 2015) or presence of additional artifact sensors (Jorge et al., 2015a) present opportunities for denoising that should be taken advantage of wherever available.

### Brain parcellation

We used the Freesurfer toolbox to process the T1-weighted images (recon-all, v6.0.0 http://surfer.nmr.mgh.harvard.edu/) in order to perform non-uniformity and intensity correction, skull stripping and grey/white matter segmentation. The cortex was parcellated into 148 regions according to the Destrieux atlas (Destrieux et al., 2010; Fischl et al., 2004) and into 68 regions according to the Desikan(-Killiany) atlas (Desikan et al., 2006). According to the results of Farahibozorg et al. (2018), showing that the optimal size of parcellation to capture independent EEG signals contains around 70 regions we decided to use the Desikan atlas as reference.

#### Specific strategy for 64Ch-7T

Due to local signal drops in the T1-weighted images near the EEG lead convergence points (Jorge et al., 2015b) we were not able to run the Freesurfer individual segmentation (recon-all) for all subjects. In order to not lose any of the subjects and to keep consistency within the data set we coregistered all fMRI images to the MNI template. We used Freesurfer to extract the surfaces of the MNI template. In order to account for subject specific variances, we dilated the atlas images by 3 voxels (using https://github.com/mattcieslak/easy_lausanne). The transformation of the segmented MNI template to fMRI was calculated by coregistering the fMRI images to the T1 image using FSL-FLIRT (6.0.2) (Jenkinson et al., 2002) with boundary-based registration using white matter masks obtained by segmentation with ANTS (version 2.2.0) (Avants et al., 2011). The T1 was coregistered to the MNI template using FSL-FNIRT (6.0.2) (Jenkinson and Smith, 2001).

##### fMRI processing

Slice timing correction was applied to the fMRI timeseries (for the Ch64-3T and Ch256-3T datasets only). This was followed by spatial realignment using the SPM12 toolbox (Ch64-1.5T/Ch64-3T: revision 6906; Ch256-3T/Ch64-7T revision 7475; http://www.fil.ion.ucl.ac.uk/spm/software/spm12). The T1 images of each subject and the Desikan/Destrieux atlas (already in subject space, as described above) were coregistered to the fMRI images (FSL-FLIRT 6.0.2). We extracted signals of no interest such as the average signals of cerebrospinal fluid (CSF) and white matter from manually defined regions of interest (ROI, 5mm sphere, Marsbar Toolbox 0.44, http://marsbar.sourceforge.net) and regressed out of the BOLD timeseries along with 6 rotation, translation motion parameters and global gray matter signal (Wirsich et al., 2017). Then we bandpass-filtered the timeseries at 0.009-0.08 Hz (Power et al., 2014). Like in Wirsich et al. (2020a, 2017), we scrubbed the data using frame wise displacement (threshold 0.5mm, by excluding the super-threshold timeframes) as defined by Power et al. (2012).

#### Specific strategy for 64Ch-7T

Data was also B0-unwarped before spatial alignment, using FSL-TOPUP (6.0.2) (Andersson et al., 2003), based on the reverse-encoding reference acquisition, to mitigate the more accentuated image distortions present at 7T (Jorge et al., 2018). Given the short TR of 1s no slice timing correction was carried out (Smith et al., 2013).

#### Specific strategy for 64Ch-1.5T

In order to stick to the original processing of (Deligianni et al., 2014) no slice timing correction was carried out. We note that as shown by Wu et al. (2011) and Shirer et al. (2015), slice timing correction has minimal to no effect on brain connectivity (in terms of test-retest reliability, signal to noise ratio and group separability).

### fMRI connectivity measures

Average timeseries of each region was then used to calculate FC_fMRI_ by taking the pairwise Pearson correlation of each regions’ cleaned timecourse (see schema Fig 1). The final connectivity matrix was constructed by the unthresholded values of the Pearson correlation.

### EEG processing

EEG data was preprocessed individually for the different setups:

#### 64Ch-1.5T

EEG was corrected for the scanner gradient artifact using template subtraction, adaptive noise cancellation and downsampling to 250Hz (Allen et al., 2000) followed by pulse-related artifact template subtraction (Allen et al., 1998). Then ICA-based denoising (for removal of gradient and pulse artifact residuals, eye-blinks, muscle artifacts) using the Brain Vision Analyzer 2 software (Brain Products, Gilching, Germany) was carried out.

#### 64Ch-3T

EEG was corrected for the scanner gradient artifact using template subtraction, adaptive noise cancellation followed by lowpass filtering at 75Hz, downsampling to 250Hz (Allen et al., 2000). Then pulse-related artifact template subtraction (Allen et al., 1998) using EEGlab v.7 (http://sccn.ucsd.edu/eeglab) and the FMRIB plug-in (https://fsl.fmrib.ox.ac.uk/eeglab/fmribplugin/) was carried out.

#### 256Ch-3T

EEG was corrected for the scanner gradient artifact using template subtraction with optimal basis set and adaptive noise cancellation (Allen et al., 2000; Niazy et al., 2005), followed by pulse-related artifact template subtraction (Allen et al., 1998) using in-house code Matlab code for ballistocardiogram peak detection as described in (Iannotti et al., 2015). Electrodes placed on the cheeks and in the face were excluded form data analysis resulting in final 204 used electrodes. This was followed by manual ICA-based denoising (for removal of gradient and pulse artifact residuals, eye-blinks, muscle artifacts, infoMax, runICA-function EEGLab revision 1.29 (Bell and Sejnowski, 1995; Delorme and Makeig, 2004))

#### 64Ch-7T

EEG data pre-processing included the following steps: gradient artifact correction using template substraction (as described in (Jorge et al., 2015a)), bad channel interpolation (1–4 channels per subject), temporal band-pass filtering (1–70 Hz), pulse-related artifact correction (using a k-means clustering-based approach validated in (Jorge et al., 2019) in line with (Gonçalves et al., 2007)), downsampling to 500 Hz, motion artifact correction (offline multi-channel recursive least-squares regression, using the motion sensor signals, as described in (Jorge et al., 2015a)), and manual ICA-based denoising (for removal of e.g. gradient and pulse artifact residuals, eye-blinks, muscle artifacts, in-house ICA extended Infomax algorithm).

#### All datasets

Cleaned EEG data was analyzed with Brainstorm software (Tadel et al., 2011), which is documented and freely available under the GNU general public license (http://neuroimage.usc.edu/brainstorm, 64Ch-1.5T and 64Ch-3T data set: version 10th August 2017 as according to (Wirsich et al., 2020b), 256Ch-3T and 64Ch-3T data set: version 15th January 2019). Data was bandpass-filtered at 0.3-70 Hz (64Ch-1.5T at 0.5-70Hz, 64Ch-7T at 1-70Hz). Data was segmented according to one TR or as a multiple TRs of the fMRI acquisition (64Ch-1.5: 2160ms, 64Ch-3T: 2000ms, 256Ch-3T: 1990ms, 64Ch-7T: sliding window of 4000ms with 1000ms (1TR) steps).

In order to minimize effect of head motion EEG epochs containing motion were semi-automatically detected if the signal in any channel exceeded the mean channel timecourse by 4 standard deviations. Then the whole timecourse was also visually inspected to exclude all motion segments from further analysis (Wirsich et al., 2020a, 2017). Electrode positions and T1 were coregistered by manually aligning the electrode positions onto the electrode artifacts visible in the T1 image. A forward model of the skull was calculated based on the individual T1 image of each subject using the OpenMEEG BEM model, (Gramfort et al., 2010; Kybic et al., 2005). The EEG signal was projected into source space (15000 solution points on the cortical surface) using the Tikhonov-regularized minimum norm (Baillet et al., 2001) with the Tikhonov parameter set to 10% (Ch64-1.5T/Ch64-3T brainstorm 2016 implementation and Ch256-3T/Ch64-7T brainstorm 2018 implementation, with default parameters: assumed SNR ratio 3.0, using current density maps, constrained sources normal to cortex with signs flipped into one direction, depth weighting 0.5/max amount 10). Finally, the source activity of each solution point was averaged in each cortical region of the Desikan and the Destrieux atlas.

### EEG connectivity measures

For each epoch the imaginary part of the coherency (iCoh, (Nolte et al., 2004)) of the source activity was calculated between each region pair (cortical regions only: Desikan atlas - 68 regions or Destrieux atlas - 148 regions) using bins of 2Hz frequency resolution (Wirsich et al., 2020a, 2017) (Brainstorm implementation, version 27-01-2019; imaginary part was corrected by the real part of the coherence 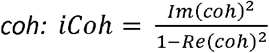 (Ewald et al., 2012), significance of each value was determined according to (Schelter et al., 2006), connections with p<0.05 were set to 0). The 2Hz bins were averaged for 5 canonical frequency bands: delta (δ 0.5-4Hz, 64Ch-7T: at 1-4Hz), theta (θ 4-8Hz), alpha (α 8-12Hz), beta (β 12-30Hz), gamma (γ 30-60Hz). The segments were then averaged for each subject to one FC_EEG_ matrix. We calculated the amplitude envelope correlation (AEC) of the signal by taking the Hilbert envelope of the concatenated epochs of each subject filtered into the canonical frequency bands both for the Destrieux and Desikan atlas. We calculated the correlation of the filtered data both without (Deligianni et al., 2014; Glomb et al., 2020) and with (Brookes et al., 2012; Hipp et al., 2012) a subsequent pairwise orthogonalization approach to attenuate crosstalk between signals (AEC_non-orthogonalized_/AEC_orthogonalized_, Brainstorm implementation; version 27-01-2019 implementation of (Hipp et al., 2012)).

### Monomodal reproducibility

We assessed the modality-specific reproducibility by determining the topographical similarity of the average connectivity matrix through calculating the monomodal inter-dataset correlation between averaged FC_fMRI_ (of each dataset across all runs and subjects). The same analysis was performed for FC_EEG_ (in each frequency band). Intra-dataset monomodal reproducibility was assessed by splitting up each dataset into two halves and calculating the correlation of the average connectome between both halves.

### Statistical analyses

To analyze the impact of group averages, we used a permutation approach that randomizes the labels of the variable of interest in order to define a p-value (5000 random permutations). In the case of statistical assessment of the 64ch-7T dataset with size of 9 subjects only label 512 permutations exist, in which case we tested for all 512 permutations to define a p-value. We report all raw p-values alongside an explicit Bonferroni threshold in case of multiple comparisons.

### The crossmodal correlation between EEG and fMRI

We then assessed the crossmodal correlation between FC_fMRI_ and FC_EEG_ for each EEG frequency band across different configurations (brain atlas, EEG connectivity measure). Effects on the crossmodal FC_EEG_-FC_fMRI_ correlation due to EEG frequency bands (δ, θ, α, β, γ), atlas choice (Desikan vs. Destrieux), EEG connectivity measure (iCoh, AEC_non-orthogonalized_, AEC_orthogonalized_) were assessed on the average connectome of each dataset (permutation test with 5000 iterations or, 512 permutations for the 64Ch-7T dataset, testing the effects against average connectomes with switched labels at the individual level, in order to be able to compare the 2278 connections of the Desikan atlas to the 11026 connections of the Destrieux atlas we randomly drew 2278 out of the 11026 Destrieux connections for each iteration. We tested if this random sampling introduces a bias to the measured crossmodal correlation by comparing 5000 (or 512 permutations for the 64Ch-7T dataset) draws of 2278 connection to the crossmodal correlation of all connections. We observed that the absolute difference was r_diff-sampling_<|0.0004|. We considered this value negligible in order to test for significant atlas differences of the order r_diff-atlas_~0.01).

To assess the effect of small sample sizes and short recording length, we cut all datasets to a recording length of the first 4min58.5s (according to the recording length in dataset 256Ch-3T). Significance was tested by randomly switching labels between 4min58.5s and full-length data (5000 iterations, or 512 permutations for the 64Ch-7T dataset). Then we took only the first nine subjects of each dataset to calculate the connectome average of EEG and fMRI (according to the number of subjects in dataset 64Ch-7T). Significance was defined by comparing the average crossmodal correlation of all subjects against the average 9 randomly sampled subjects from the group (5000 iterations).

### Spatial characterization of the crossmodal correlation

Focusing on the multimodal connectomes in the Desikan atlas averaged over all datasets, we assessed the relative contribution of each connection to the correlation between FC_EEG_ and FC_fMRI_ according to Colclough et al. (2016). The relative contribution c of each connection i is given by: 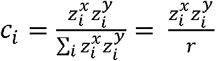 with 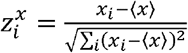 and 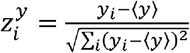 given the Pearson correlation coefficient of two vectors x and 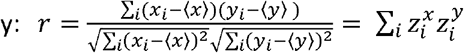. The resulting spatial contribution matrix was then correlated across the different datasets to assess the reproducibly of this spatial contribution to the crossmodal relationship. To classify the results we mapped the 68 regions of the Desikan atlas to 7 canonical ICNs (VIS: Visual, SM: Somatomotor, DA: Dorsal Attention, VA: Ventral Attention, L: Limbic, FP: Fronto-Parietal, DMN: Default Mode Network) (Amico et al., 2017; Yeo et al., 2011) and we subdivided the connections into homotopic, intrahemispheric and interhemispheric connections. We tested if the spatial contribution to the crossmodal correlation was more prominent for each ICN, for homotopic connections and for interhemispheric connections (one-sided ttest). Finally, we tested if the magnitude of the crossmodal correlation was increased for each ICN, for homotopic connections and for intrahemispheric connections (one-sided ttest).

### Reproducibility of the crossmodal correlation from single subject to the average connectome

Besides taking the average multimodal connectome of each dataset we also calculated the correlation between EEG and fMRI connectomes in each subject. Using a two-way ANOVA we tested if the crossmodal correlation differs in terms of the dataset and EEG frequency band or the interaction between dataset and frequency band (Matlab function ANOVAN, p<0.05). To identify any possible effects of a specific dataset or frequency band on the crossmodal correlation, we used a Tukey Posthoc test (Matlab function multcompare, p<0.05 Bonferroni corrected). In order to exclude a large effect of motion on the crossmodal relationship we correlated the average framewise displacement of each subject to the individual crossmodal relationship (see SI section impact of movement). Additionally, to limiting the group average to 9 subjects, in order to better understand the relationship between single subject and dataset averaged estimates we aimed to define the number of subjects needed for a reliable averaged connectome. To do so now and randomly selected 1,2,…,n subjects (5000 iterations each step) and averaged the EEG and fMRI connectomes of the selected subjects. Then we calculated the correlation between FC_EEG_ and FC_fMRI_ for each of the subject steps. We repeated this approach for the combined dataset sampling an average correlation between the averaged EEG and fMRI connectome from n randomly drawn subjects. To determine how many subjects are needed for a stable average connectome we took the average crossmodal correlation over all subjects of each frequency band as reference correlation. We then compared the value of 1% of reference correlation to the change rate when adding one random subject (5000 iterations). The crossmodal correlation was considered stable when the change rate did not differ more than 1% of the total crossmodal reference correlation.

## Results

### Monomodal reproducibility between datasets

We measured monomodal reproducibility (topographical similarity) by taking correlations of connectivities of each modality. Between datasets, monomodal connectivity matrices were all correlated for both modalities (Table 1, Table 2). 7T fMRI data correlated less with the other datasets (Table 1), EEG correlations were lower than correlations between fMRI (Table 1, Table 2). In terms of inter-dataset connectome correlation, delta, theta and beta band connectomes were the most correlated across datasets (r>0.65, see Table 2). Unlike otherwise stated, the results of this section are all derived from the group averaged connectomes of each dataset.

**Table 1:**
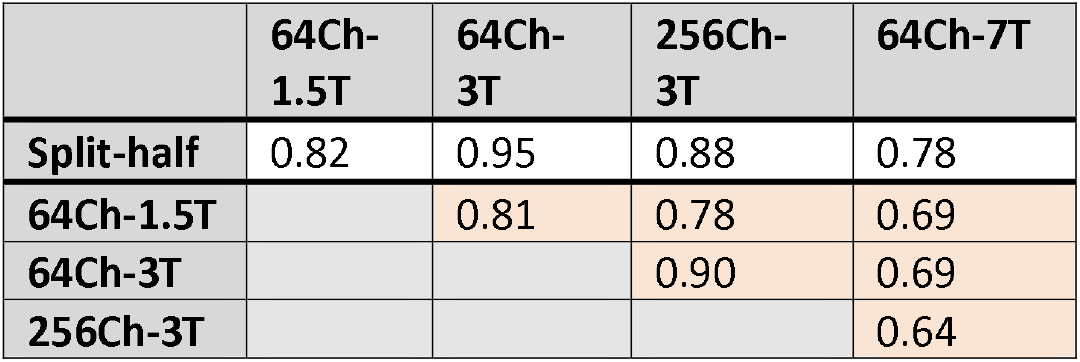
Inter- and intra-dataset correlation of FC_fMRI_.; The first row shows the intra-dataset correlation of the dataset’s split-half averaged fMRI connectome. The orange cells show the inter-dataset correlation of dataset average fMRI connectome (Desikan atlas) between the different datasets

|  | 64Ch-1.5T | 64Ch-3T | 256Ch-3T | 64Ch-7T |
| --- | --- | --- | --- | --- |
| **Split-half** | 0.82 | 0.95 | 0.88 | 0.78 |
| **64Ch-1.5T** |  | 0.81 | 0.78 | 0.69 |
| **64Ch-3T** |  |  | 0.90 | 0.69 |
| **256Ch-3T** |  |  |  | 0.64 |

**Table 2:**
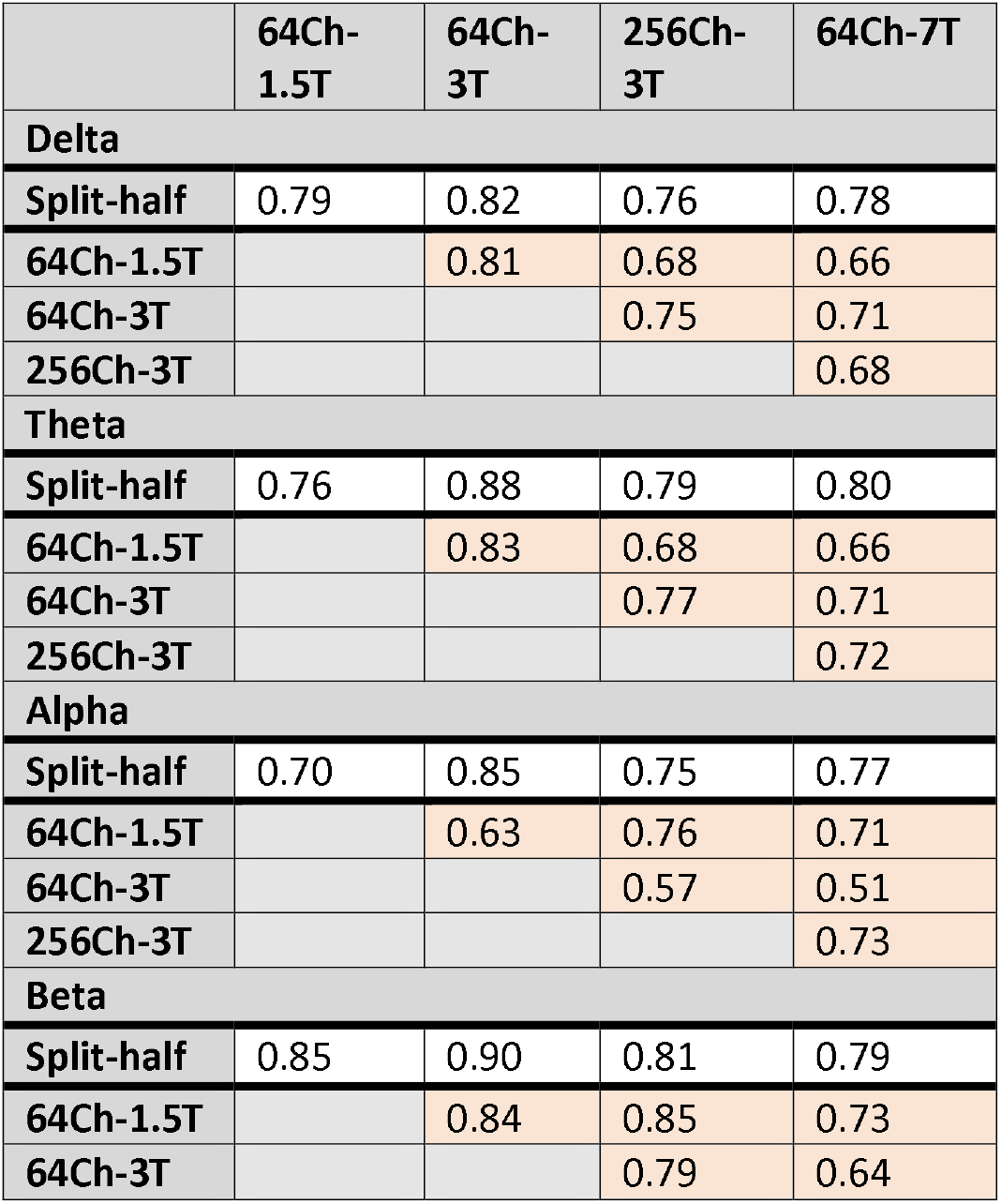
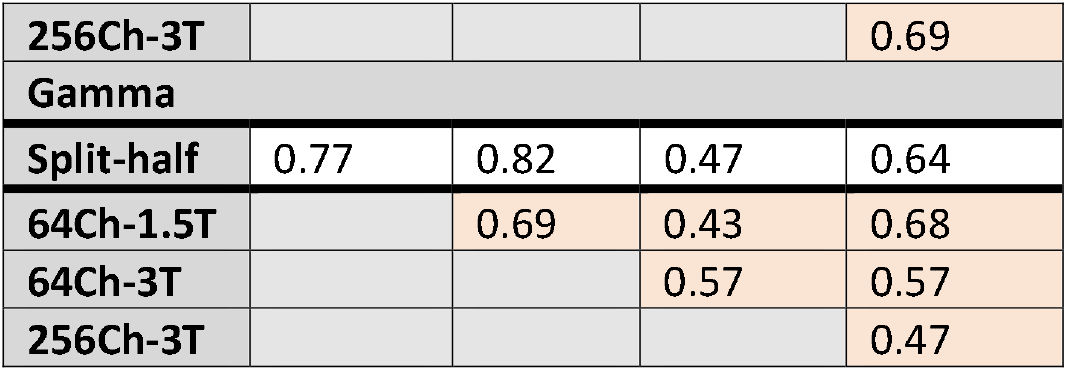
Inter- and intra-dataset correlation of FC_EEG_. The first row shows of each frequency-band the intra-dataset correlation of the dataset’s split-half averaged EEG connectome. The orange cells show the inter-dataset correlation of dataset averaged EEG connectome (Desikan atlas, imaginary part of the coherency) between the different datasets.

### Crossmodal correlation of group averaged EEG and fMRI connectomes

Correlations between EEG and fMRI were highest for the beta band, while gamma and delta band were the most variable across datasets. A grand average across all datasets (72 subjects) resulted in the highest correlation between EEG and fMRI for all bands (Fig 2a). While the grand average FC_fMRI_-FC_EEG-β_ correlation was significantly higher than all the other bands (p<0.0002, 5000 permutations, see SI Table 3), the group-averaged FC_fMRI_-FC_EEG-α_ correlation was significantly increased only when compared to FC_fMRI_-FC_EEG-γ_ (only for 64Ch-3T dataset: p=0.0002 and 256-3T dataset: p=0.0004, 5000 permutations, see SI Table 3).

**Fig 1:**
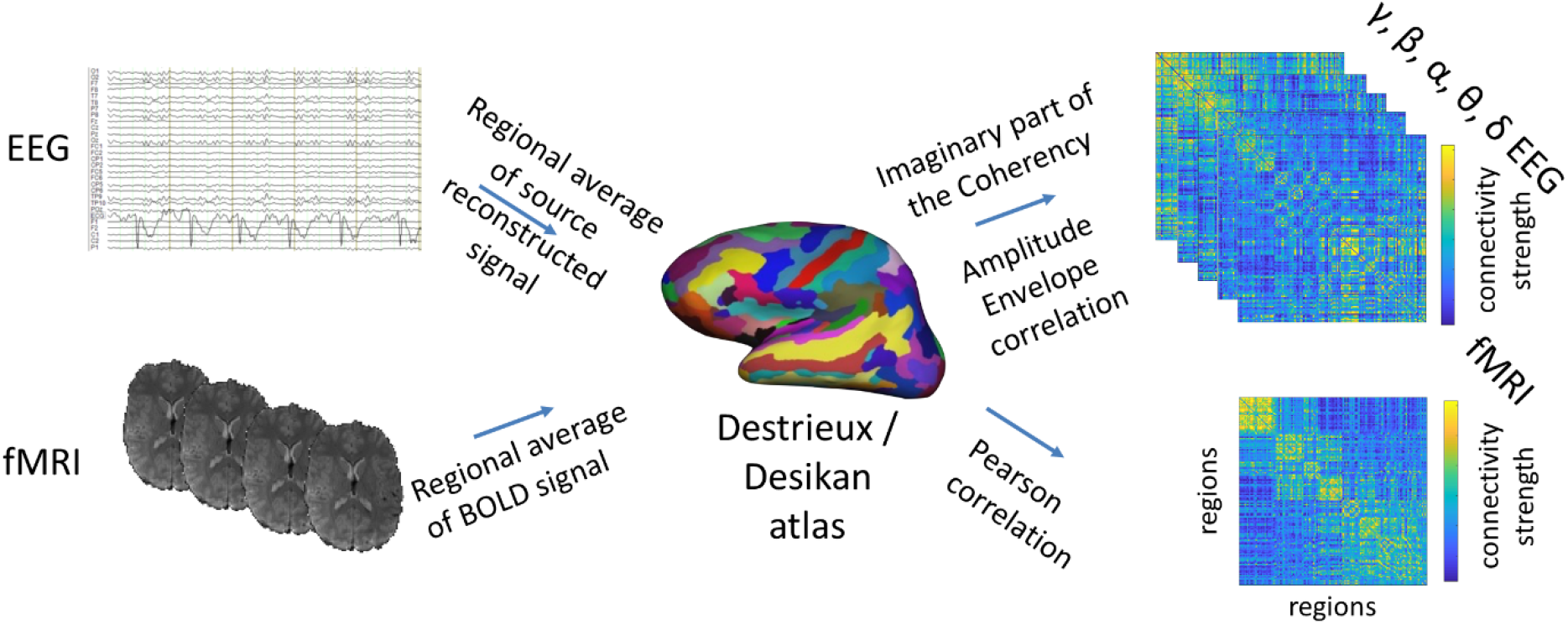
Overview on the construction of EEG and fMRI connectomes. EEG and fMRI data were parcellated into the 148 cortical regions of the Destrieux atlas (and 68 regions of the Desikan atlas, coregistered to each subject’s individual T1) as follows: For fMRI, the BOLD signal timecourse was averaged over the voxels in each region for each subject. The Pearson correlation of the region averaged fMRI-BOLD timecourse was calculated to build a function connectivity matrix /connectome (FC_fMRI_). For the EEG, the signal of each sensor was source reconstructed to the cortical surface (15000 solution points) using the Tikhonov-regularized minimum norm. Then, the timecourses of the solution points were averaged per cortical region. The imaginary part of the coherency (iCoh) or envelope amplitudes correlation (AEC, orthogonalized and non-orthogonalized) of averaged EEG source signals were used to calculate FC_EEG_ for each subject (Figure adapted from (Wirsich et al., 2020a)). Please refer to the methods for a detailed description of each step.

**Fig 2:**
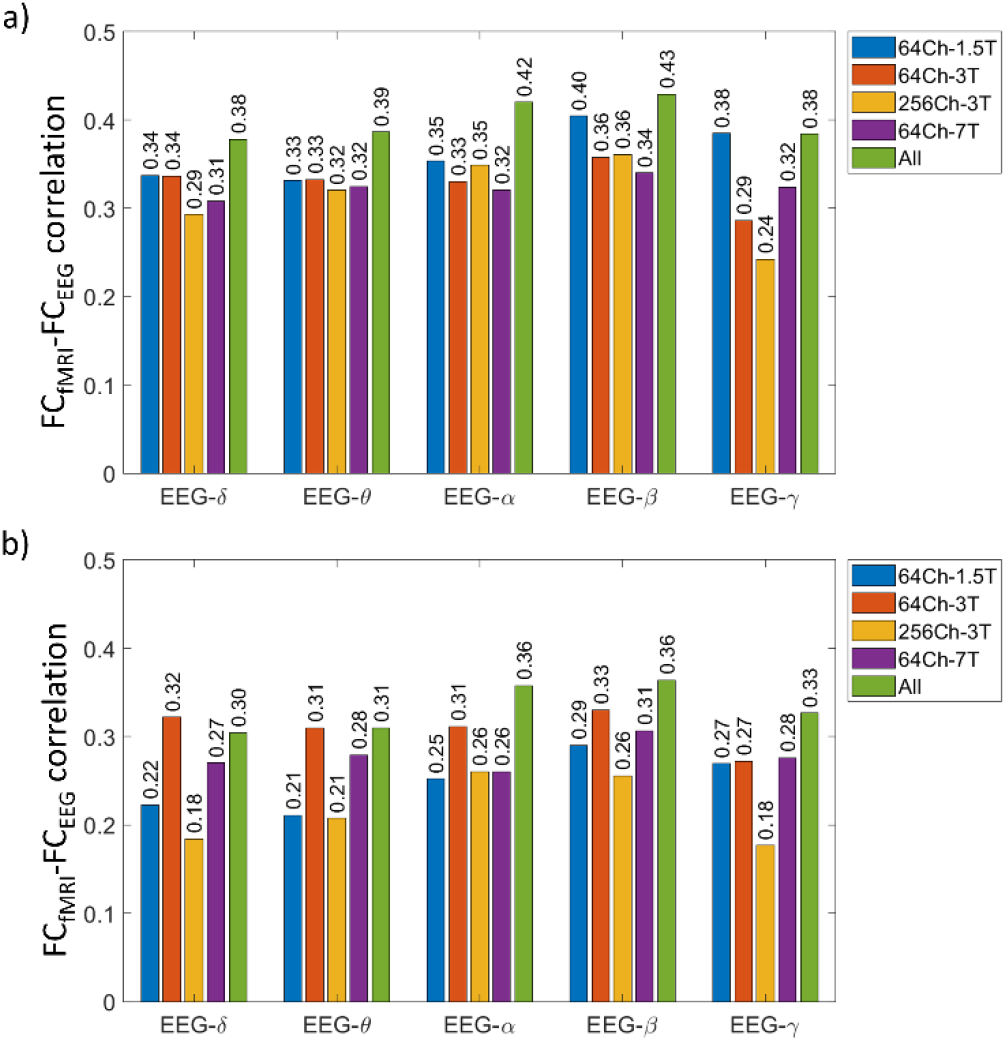
a) Crossmodal correlation dataset averaged EEG and fMRI connectomes using the Desikan atlas (FC_EEG_ measure: imaginary part of the coherency); b) Crossmodal correlation dataset averaged EEG and fMRI connectomes using the Destrieux atlas (FC_EEG_ measure: imaginary part of the coherency).

### Crossmodal correlation of group averaged EEG and fMRI connectomes using an alternative atlas and alternative EEG connectivity measures

In this section we compared the above used atlas (Desikan) and EEG connectivity measures (iCoh) to alternatives. Taking an atlas with higher resolution resulted in a reduction of crossmodal FC_fMRI_-FC_EEG_ correlation grand average over all datasets (Fig 2b, p<0.0002 for all frequency bands, 5000 permutations, see SI Table 4, a subset of the Destrieux connections were randomly chosen for each iteration to match the number of connections of the Desikan atlas: see methods). When using AEC_non-orthogonal_ as FC_EEG_ connectivity measure crossmodal correlation increased as compared to taking iCoh (Fig 3b, for FC_EEG-γ_ only, p<0.0002, 5000 permutations, SI Table 5), the orthogonalization approach (AEC_orthogonal_) resulted in lower correlation compared to the iCoh (Fig 3b, all EEG frequency bands, p<0.0002, 5000 permutations, see SI Table 5). We did not find any significant correlation between FC_fMRI_ and FC_EEG-γ_ in the 256Ch-3T dataset.

**Fig 3:**
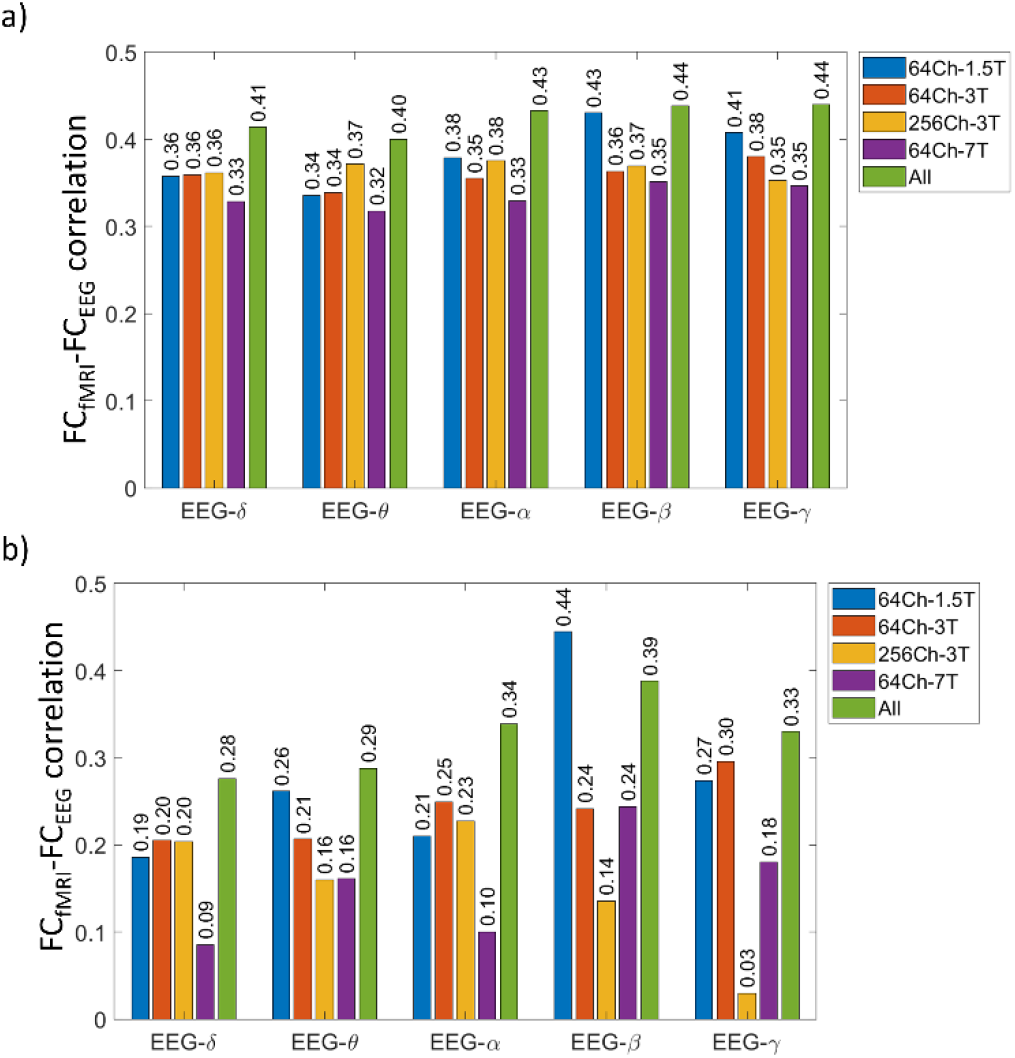
Crossmodal correlation dataset averaged EEG and fMRI connectomes using Amplitude envelope correlation (AEC) to derive FC_EEG_ a) non-orthogonalized AEC and b) orthogonalized AEC (results for Desikan atlas)

### Effect of number of subjects and length of resting-state recording on the crossmodal correlation

Next connectomes were cut down to the session length of the shortest dataset (4min58.5s, Fig 4a) and subject averages were calculated using the same number of subjects (9 subjects, Fig 4b). The crossmodal correlation averaged over all datasets is not significantly different when the sessions are cut down to the first 4min58.5s (p>0.07 for all frequency bands, 5000 permutations, SI Table 6). Note that though we did not find any significant differences for group averaged connectomes, we observed significant differences individual crossmodal correlation when limiting the 30 minutes of the 64Ch-3T dataset to 4min58.5s (one-sided ttest, significant for FC_fMRI_-FC_EEG-δ_, FC_fMRI_-FC_EEG-θ_ and FC_fMRI_-FC_EEG-α_, p<0.0003, SI Table 7). When taking the average connectivity of only the first 9 subjects the crossmodal correlation between FC_fMRI_ and FC_EEG-γ_ decreases significantly (p=0.0006, 5000 permutations, SI Table 6).

**Fig 4:**
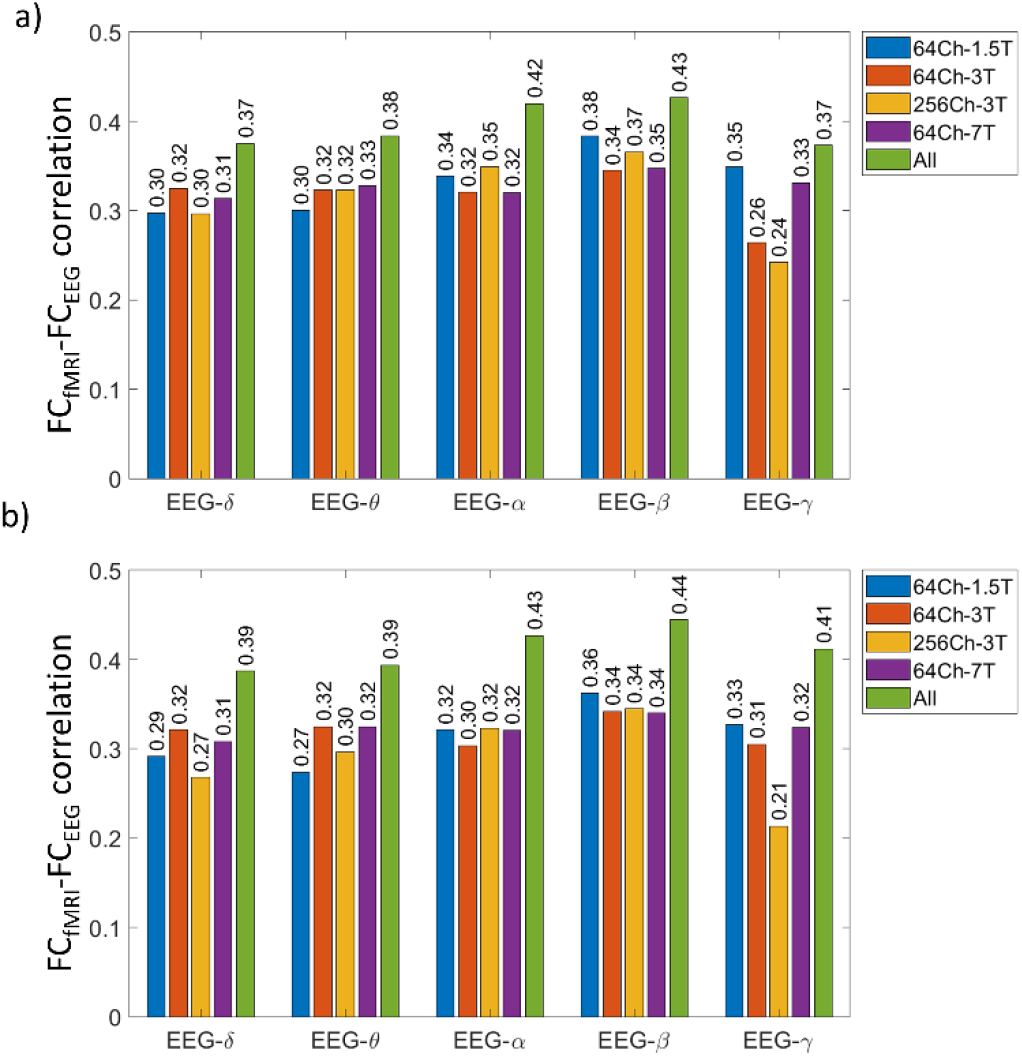
Crossmodal correlation between dataset averaged EEG and fMRI connectomes limiting all datasets to: a) the first 4min58.5s of each subject’s session (according to the session length of the 256Ch-3T dataset) and b) the first 9 subjects of each dataset (according to the number of subjects of the 64Ch-7T dataset).

### Spatial characterization of differences and ICN crossmodal correlation

We defined the connections that contribute most to the FC_fMRI_-FC_EEG_ correlation using the connectomes averaged over all datasets (Colclough et al., 2016). Table 3 shows that the topography this spatial contribution is correlated between all datasets. The visual network contributes the most to the crossmodal correlation as well as the homotopic connections (Fig 5). Connections of the visual network contributed significantly more to the crossmodal FC_fMRI_-FC_EEG_ correlation as compared to inter-ICN connections for all frequency bands (t-test spatial contribution Visual>interICN: FC_fMRI_-FC_EEG-δ_/FC_EEG-θ_/FC_EEG-α_/FC_EEG-β_/FC_EEG-γ_; p=3.2*10^−47^/1.8*10^−34^/p=5.1*10^−69^/p=4.1*10^−53^/1.5*10^−81^, Bonferroni correction threshold for 5 frequencies and 7 ICNs is defined at p=0.05/35=0.0014). Connections of the limbic network contributed significantly more to the FC_fMRI_-FC_EEG_ correlation as compared to inter-ICN connections for all frequency bands except EEG-γ (t-test spatial contribution Limbic>inter-ICN: FC_fMRI_-FC_EEG-δ_/FC_EEG-θ_/FC_EEG-α_/FC_EEG-β_; p=5.4*10^−06^/4.3*10^−06^/4.6*10^−05^/3.2*10^−05^, Bonferroni correction threshold for 5 frequencies and 7 ICNs is defined at p=0.05/35=0.0014).

**Table 3:** Inter- and intra-dataset correlation of spatial region-specific contributions to the averaged FC_fMRI_-FC_EEG_ crossmodal correlation. The first row of each frequency-band shows the intra-dataset correlation of the dataset’s split half averaged EEG-fMRI connectomes. The orange cells show the inter-dataset correlation of the dataset averaged EEG-fMRI connectome (Desikan atlas, imaginary part of the coherency).

| Spatial contribution | 64Ch-1.5T | 64Ch-3T | 256Ch-3T | 64Ch-7T |
| --- | --- | --- | --- | --- |
| **Delta** |  |  |  |  |
| Split-half | 0.72 | 0.85 | 0.76 | 0.79 |
| 64Ch-1.5T |  | 0.76 | 0.62 | 0.64 |
| 64Ch-3T |  |  | 0.76 | 0.64 |
| 256Ch-3T |  |  |  | 0.61 |
| **Theta** |  |  |  |  |
| Split-half | 0.69 | 0.87 | 0.81 | 0.80 |
| 64Ch-1.5T |  | 0.76 | 0.65 | 0.61 |
| 64Ch-3T |  |  | 0.78 | 0.62 |
| 256Ch-3T |  |  |  | 0.64 |
| **Alpha** |  |  |  |  |
| Split-half | 0.72 | 0.84 | 0.80 | 0.79 |
| 64Ch-1.5T |  | 0.56 | 0.75 | 0.76 |
| 64Ch-3T |  |  | 0.62 | 0.50 |
| 256Ch-3T |  |  |  | 0.69 |

| Beta |  |  |  |  |
| --- | --- | --- | --- | --- |
| Split-half | 0.82 | 0.88 | 0.86 | 0.81 |
| 64Ch-1.5T |  | 0.81 | 0.80 | 0.74 |
| 64Ch-3T |  |  | 0.84 | 0.65 |
| 256Ch-3T |  |  |  | 0.66 |
| Gamma |  |  |  |  |
| Split-half | 0.80 | 0.86 | 0.65 | 0.75 |
| 64Ch-1.5T |  | 0.79 | 0.66 | 0.70 |
| 64Ch-3T |  |  | 0.71 | 0.66 |
| 256Ch-3T |  |  |  | 0.61 |

**Fig 5:**
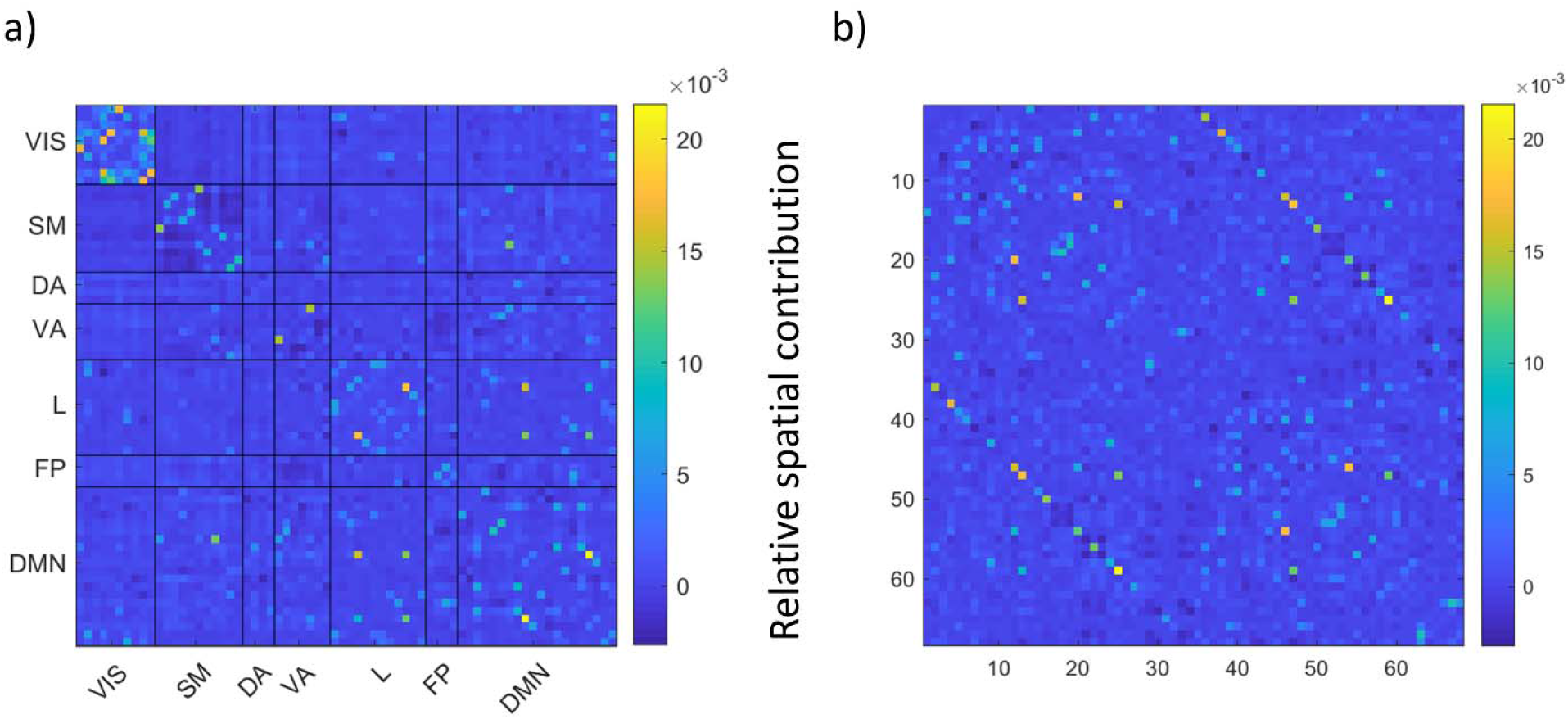
Relative spatial contribution (see methods) of each connection to the crossmodal correlation between FC_EEG-β_ and FC_fMRI_ based on the average over all 72 subjects. (a) Spatial contribution is ordered according to the 7 canonical ICNs (Yeo et al. 2010). The Visual Network and the Limibic Network contributed the most to the crossmodal relationship. (b) Spatial contribution is ordered according to the two hemispheres. Off-diagonals highlight the homotopic connections that contributed the most to the crossmodal correlation. Abbreviations: VIS: Visual, SM: somatomotor, DA: dorsal attention, VA: ventral attention, L: Limbic, FP: Fronto parietal, DMN: default mode network

Homotopic connections contributed significantly more to the FC_fMRI_-FC_EEG_ correlation as compared to the rest of the connectome for all bands (t-test spatial contribution homotopic>other connections: FC_fMRI_-FC_EEG-δ_/FC_EEG-θ_/FC_EEG-α_/FC_EEG-β_/FC_EEG-γ_: p=2.1*10^−52^/8.710*10^−50^/4.3*10^−52^/p=1.2*10^−55^/9.6*10^−56^, Bonferroni correction threshold for 5 frequencies is defined at p=0.05/5=0.01). Additionally, intrahemispheric connections contributed significantly more than interhemispheric connections to the crossmodal relationship (t-test spatial contribution intrahemispheric>interhemispheric: FC_fMRI_-FC_EEG-δ_/FC_EEG-θ_/FC_EEG-α_/FC_EEG-β_/FC_EEG-γ_: p=1.1*10^−13^/1.3*10^−14^/1.6*10^−16^/9.3*10^−23^/5.9*10^−28^, Bonferroni correction threshold for 5 frequencies is defined at p=0.05/5=0.01)

To follow up we explored at the crossmodal correlation of FC_fMRI_-FC_EEG_ while only selecting connections inside one ICN. When comparing the crossmodal correlation inside the different canonical ICNs to the crossmodal correlation of random in-between network connections of the same network size (randomly sampled, 5000 iterations), we observe a higher correlation in the Visual network (r_fMRI_,_EEG-α_ =0.61 (r>r_random_: p=0.028), r_fMR,EEG-γ_=0.63 (r>r_random_: p=0.007), the Limbic network (r_fMRI_,_EEG-δ_=0.56, (r>r_random_: p=0.03), r_fMRI,EEG-β_=0.59 (r>r_random_: p=0.022), r_fMRI,EEG-γ_=0.60 (r>r_random_: p=0.0034)) and the DMN (r_fMRI,EEG-γ_=0.52 (r>r_random_: p<0.0002), all significant at uncorrected threshold p<0.05, Bonferroni correction threshold is defined at p=0.05/35=0.0014).

### Reproducibility of crossmodal correlation in individual connectomes

We generally observed the same distribution of FC_EEG_-FC_fMRI_ correlation in individual as compared to the dataset average such as high crossmodal correlation for FC_EEG-β_ and low crossmodal correlation for FC_EEG-γ_ (Table 4). Lowest correlation was observed in the 256Ch-3T dataset.

**Table 4:**
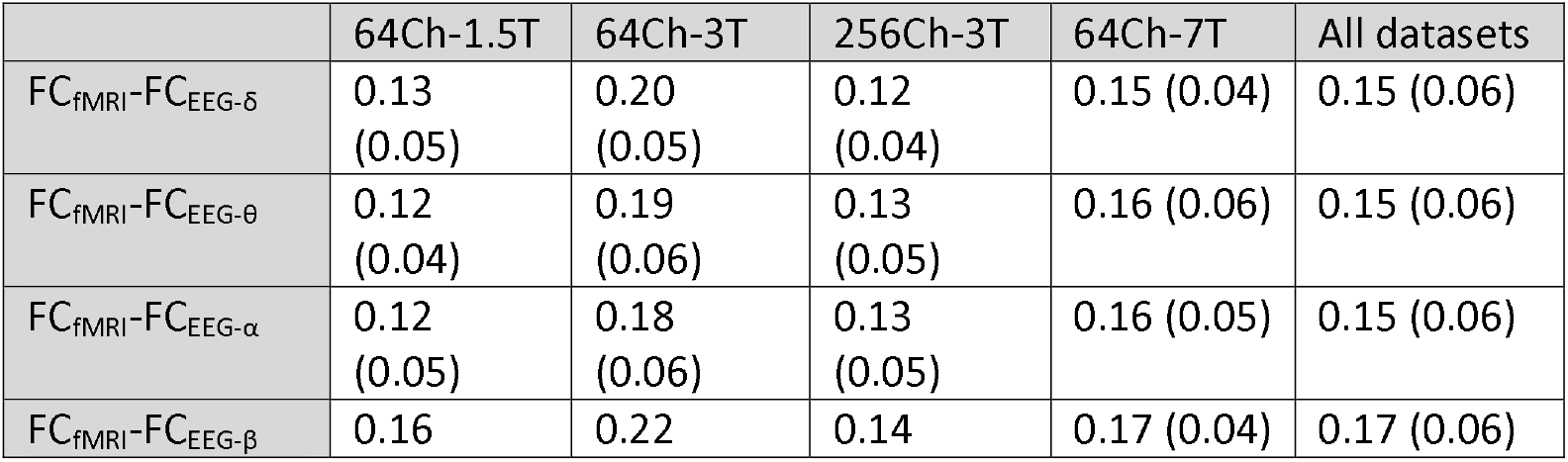
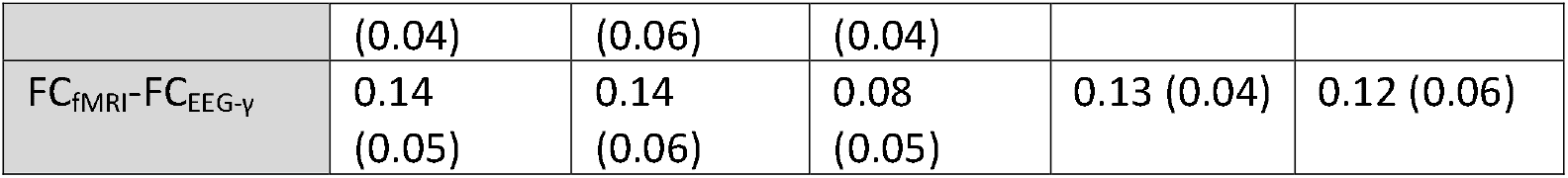
Average crossmodal correlation between EEG and fMRI connectomes of individuals across each dataset and frequency band (standard deviation in brackets)

A 2-way ANOVA of the individual subject crossmodal correlation revealed a significant main effect of datasets F(3, 71)=36.61, p=1.6*10^−20^; and a significant main Effect of EEG Frequency bands F(4, 71)=6.85, p=2.6*10^−5^. The interaction term dataset*band was not significant (F(12, 71)=1.32, p=0.21, significance threshold p<0.05). Tukey Posthoc t-tests on the main effect of the datasets established the following order of correlation magnitude: 64Ch-3T>64Ch-7T>64Ch-1.5T>256Ch-3T (64Ch-3T > 64Ch-1.5T (p<0.0001), 64Ch-3T>64Ch-7T (p=0.0031), 64Ch-3T>256Ch-3T (p<0.0001); 64Ch-7T>256Ch-3T (p=0.003)). Tukey Posthoc t-tests on the main effect of EEG frequency bands revealed that FC_fMRI_-FC_EEG-γ_ correlation was significantly smaller than FC_fMRI_-FC_EEG-β_ (p<0.0001) and FC_fMRI_-FC_EEG-θ_ correlation (p=0.027).

FC_EEG-β_ correlates the best with FC_fMRI_ for all datasets (Fig 6, Fig 7). On the contrary FC_EEG-γ_-FC_fMRI_ was dependent on the data set showing the 2^nd^ strongest correlation for the 64Ch-1.5T dataset and being by far the lowest correlation for the 64Ch-3T and 256Ch-3T dataset (Fig 7). Independent of the magnitude of the crossmodal correlation of each EEG frequency band, the crossmodal correlation was observed to be stable for each dataset when averaging 7-12 subjects (stable = adding 1 subject did not change the correlation more than 1% of the total crossmodal correlation when averaging connectomes over 72 subjects, SI Table 2). Combining all dataset to one average connectome maximized the crossmodal relationship for between FC_fMRI_ and FC_EEG-δ_, FC_EEG-θ_, FC_EEG-α_ and FC_EEG-β_ but not for FC_EEG-γ_. The 64Ch-1.5T dataset has an exceptionally high crossmodal correlation between FC_EEG-γ_ and FC_fMRI_. Consequently the correlation is not increased by taking the average of all 72 subjects (and effect clearly observed for all other bands Fig 2a: 64Ch-1.5T dataset: r(FC_EEG-γ_, FC_fMRI_)=0.38 vs. all datasets: r(FC_EEG-γ_, FC_fMRI_)=0.39).

**Fig 6:**
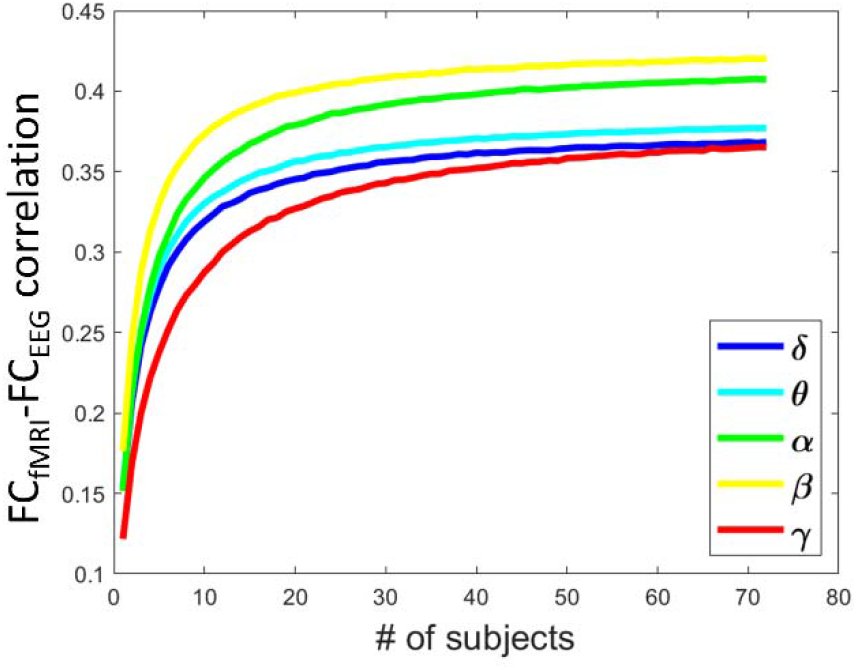
Subjects were randomly sampled from all datasets taking 1…n subjects (5000 iterations) then the crossmodal correlation between the EEG and fMRI connectome of each frequency band was calculated, the crossmodal correlation does not change more than 1% of the maximum value after averaging around 7-12 subjects (see SI Table 2). For maximum correlation see also Fig 2 (all datasets).

**Fig 7:**
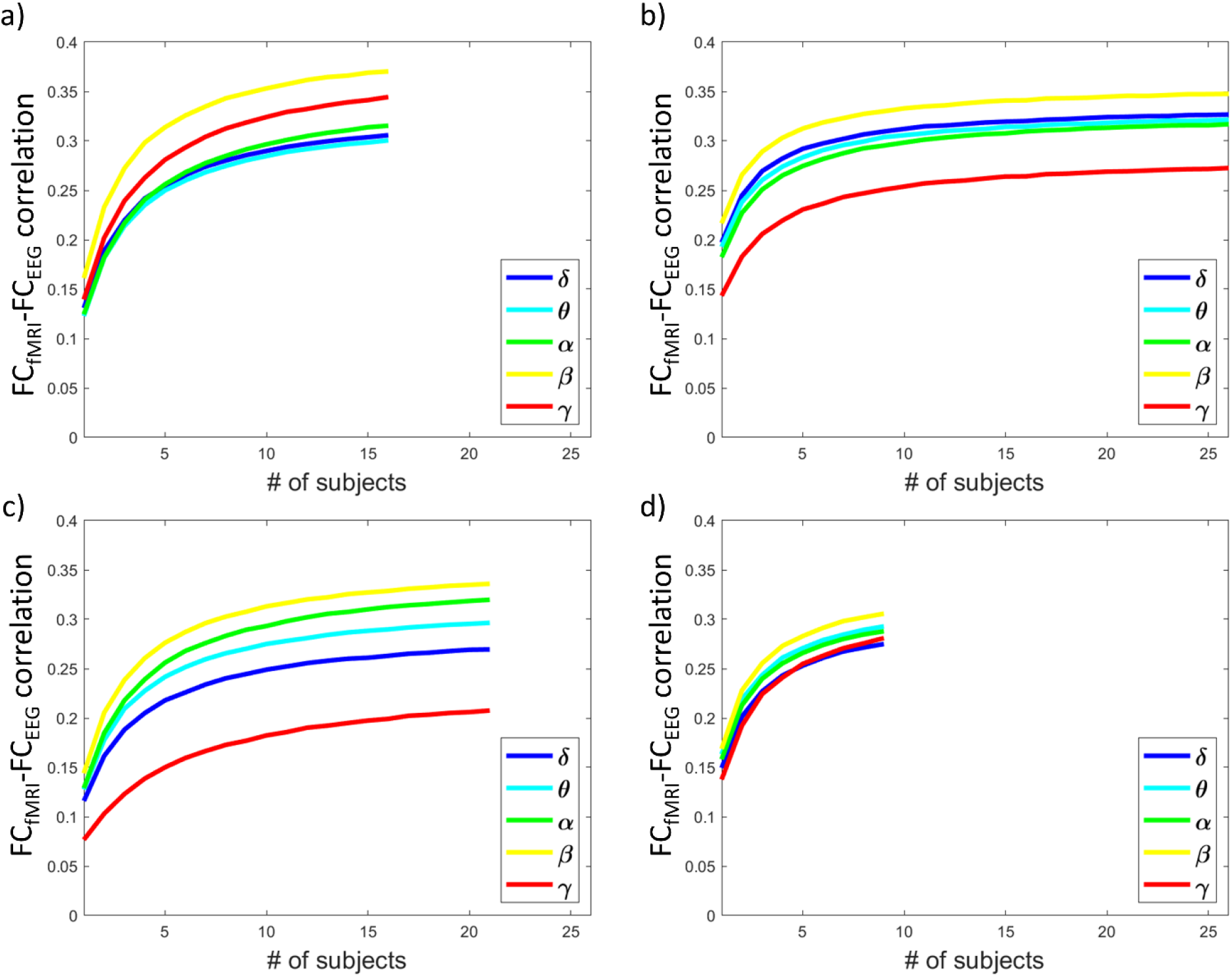
Subjects were randomly sampled for each dataset separately taking 1…n subjects (5000 iterations) then the crossmodal correlation between FC_fMRI_ and the FC_EEG_ of each frequency band was calculated, correlation does not change more than 1% of the maximum value after averaging around 7-12 subjects (SI Table 2). a) 64Ch-1.5T b) 64Ch-3T c) 256Ch-3T d) 64Ch-7T. While the crossmodal correlation is maximum for FC_fMRI_-FC_EEG-β_ FC_fMRI_-FC_EEG-γ_ is lowest for all datasets except the 64Ch-1.5 dataset (a). For the average crossmodal correlation across each dataset see also Fig 2.

## Discussion

This study showed that the crossmodal correlation between EEG and fMRI connectivity can be simultaneously recorded with high reproducibility from a variety of experimental setups and designs, notably different MR magnetic fields and EEG electrode configurations (monomodal connectivity correlation for EEG and fMRI r~0.5-0.9 between all datasets and crossmodal connectivity correlation r~0.3-0.4 across all datasets). Of special note, we demonstrated for the first time that concurrent EEG and fMRI connectomes derived from 7T show the same monomodal and crossmodal correlations as compared to data from 1.5T and 3T. From an EEG-frequency point of view the crossmodal correlation was highest for FC_fMRI_-FC_EEG-β_, while from a spatial point of view the visual network and homotopic connections contributed the most to the crossmodal correlation. When averaging subjects across all datasets, the correlation reaches a stable value from 7-12 subjects (see Fig 6, SI Table 2). From a single-subject point of view, crossmodal correlation was weak (r~0.12-0.2). When comparing the correlation to the more established crossmodal relationships between FC_fMRI_ and SC_dMRI_ we note that our correlations between EEG and fMRI have the same order of magnitude for both group averaged connectomes (e.g. r=0.36 (Honey et al., 2009), r=0.34 (Goñi et al., 2014)) and single-subject connectomes (e.g. r=0.18 (Skudlarski et al., 2008), r=0.19 (van den Heuvel MP et al., 2013)).

### Frequency specific contributions

When combining all datasets, FC_EEG-α_ and FC_EEG-β_ correlated best with FC_fMRI_ (r_fMRI-EEG-α_=0.41 and r_fMRI-EEG-β_=0.43, average over all datasets, see Fig 2a). The weaker crossmodal correlation between FC_EEG-γ_/FC_EEG-δ_ and FC_fMRI_ is in line with previous findings (Tewarie et al., 2016; Wirsich et al., 2017). These results suggest that, besides band-specific SNR observed in our monomodal inter- and intra-dataset topographical similarity (which would predict the strongest crossmodal coupling between FC_fMRI_ and FC_EEG-α_ instead of FC_-EEG-β_ (Colclough et al., 2016; Marquetand et al., 2019)), frequency specific FC_fMRI_-FC_EEG_ correlation can advance the functional understanding of large-scale connectivity. Specifically, the particularly strong tie between FC_EEG-α_ / FC_EEG-β_ and FC_fMRI_ may suggest that phase synchrony in α- and β-band contribute particularly strongly to the intrinsic network organization of the brain first characterized in FC_fMRI_, paralleling conclusions of prior MEG studies (Brookes et al., 2011; Hipp et al., 2012).

This conclusion may come as a surprise, since neurophysiological investigations using intracranial studies (human or animal) have demonstrated highest correlation between local BOLD amplitudes and γ power (Logothetis et al., 2001; Nir et al., 2007). In this context, it is important keep in mind the difference between local signal amplitudes and whole-brain FC organization. Further, we note that beyond FC_EEG-α/β_ we found weaker but significant correlations between FC_fMRI_ and FC_EEG-γ_ (as well as FC_EEG-δ_) for all connectivity measures with the exception of FC_fMRI_-FC_EEG-γ_ correlation in the 256Ch-3T dataset when using orthogonalized amplitude envelope correlations (AEC) (see also section Low SNR of gamma). Additionally, in our previous work we have shown that local (visual) FC_EEG-γ_ as well as distributed FC_EEG-δ_ provide additional information (beyond FC_EEG-α_ and FC_EEG-β_) to explain structural connectivity derived from dMRI (Wirsich et al., 2017). Further, we recently demonstrated that FC_EEG-γ_ provides spatially independent information to the FC_fMRI_-FC_EEG_ relationship (Wirsich et al., 2020a). A frequency specific crossmodal relationship is further to be expected from the laminar organization of the brain linking EEG-γ activity to local laminar connectivity while lower frequencies support long range projections (Scheeringa et al., 2016; Scheeringa and Fries, 2017). To conclude, FC_EEG-γ_ meaningfully and uniquely relates to FC_fMRI_ albeit at a weaker effect size.

### Spatial contributions and ICN organization across timescales

We observed that ICNs in general and particularly the connections of the visual and the limbic network consistently contributed more than inter-ICN connections to the static crossmodal correlation across all frequency bands. Intra-ICN connections do not only have the highest connectivity but were also previously found to have the least dynamic connections in the brain (Zalesky et al., 2014). Together with our results, this suggests the existence of a crossmodal static core component present in both EEG and fMRI (Sadaghiani and Wirsich, 2020), potentially mediated by the structural core of the brain (van den Heuvel and Sporns, 2011; Wirsich et al., 2017). This idea of a crossmodal core is further strengthened by the observation of a dominance of homotopic connections to the relationship, in line with (Shen et al., 2015) showing that homotopic regions are among the least dynamic connections of FC_fMRI_. In the current study we extend these results of (Shen et al., 2015; Wirsich et al., 2020a; Zalesky et al., 2014) by showing that visual, limbic (temporo-orbitofrontal) and homotopic connections are the largest contributors to the static crossmodal FC_fMRI_-FC_EEG_ relationship. If this property would be only driven by SNR of the EEG signal, we would expect that FC_EEG-β_ performs equally well when comparing our results to the dynamic crossmodal relationship. As we demonstrated recently (Wirsich et al., 2020b) this is not the case: the dynamic crossmodal relationship is dominated by long-range intra-ICN FC_EEG-δ_, suggesting different frequency-specific crossmodal relationships between EEG and fMRI. Taken together, this suggests a static FC_fMRI_ core dominated by correlation to FC_EEG-β_ while the results of Wirsich et al. (2020b) suggest a tight link of long-range dynamic FC_EEG-δ_ linked to dynamic FC_fMRI_.

### The relationship between FC_fMRI_ and FC_EEG_ derived by imaginary part of coherency and amplitude envelope correlation

We confirmed that correlation of amplitude envelope correlations (AEC) are related to fMRI (Deligianni et al., 2014) and provide consistent estimates of connectivity (Colclough et al., 2016). While AEC_non-orthogonalized_ correlated the most to FC_fMRI_, it has been noted that this cross-measure consistency might stem from source leakage. It has been proposed that source leakage can be solved by orthogonalization of the signal pairs before calculating the AEC (Colclough et al., 2016). Colclough et al. (2016) observed a poorer (inter- and intrasubject) topographical similarity of coherence-based measures as compared to AEC. Exploring the FC_EEG_-FC_fMRI_ correlation across datasets, we observed that the imaginary part of the coherency (iCoh) had a higher crossmodal correlation than AEC_orthogonalized_ but lower crossmodal correlation when compared to AEC_non-orthogonalized_. As we observed higher correlation of AEC-based FC_EEG_ with FC_fMRI_ when orthogonalization was not applied, it might be the case that the orthogonalization eliminates true FC_EEG_ correlated to FC_fMRI_. Instead of orthogonalization, Glomb et al. (2020) proposed to filter the FC_EEG_ using structural connectivity derived by dMRI. In that sense, it could be argued that AEC_non-orthogonalized_ provides the best estimation of neural connectivity as it has been shown to be both the most reliable measure of FC_EEG_ (Colclough et al., 2016) and to have a higher crossmodal correlation to FC_fMRI_ as compared to iCoh and AEC_orthognalized_ (Fig 3). Further work is needed to better understand the contribution of veridic neuronal connectivity at zero lag (Engel et al., 1991) that is suppressed by either using orthogonalization or iCoh. As the magnitude of the crossmodal correlation of iCoh lies between the values for orthogonalized and non-orthognalized AEC, we conclude that our results provide further evidence for the overall concordance of amplitude-based and phase-based static connectivity during resting-state (Mostame and Sadaghiani, 2020; Sadaghiani and Wirsich, 2020). Beyond the measurement of undirected connectivity, EEG has been shown to be able to extract the directional connectivity of ICN organization (Coito et al., 2019). Ultimately a multimodal approach holds the promise to better estimate directional connectivity (Wei et al., 2020).

### Crossmodal correlation of high-quality EEG-fMRI connectomes at 7T

We demonstrated for the first time that simultaneous EEG-fMRI connectivity estimation can be undertaken at 7T, providing reproducible monomodal estimates of FC_fMRI_ and FC_EEG_ comparable to data at 3T and 1.5T. EEG connectomes remain generally reproducible at 64Ch-7T when compared to the other datasets (inter-dataset correlation of FC_EEG_: r>0.51 vs. r>0.43 for all other datasets, Table 2). This is despite the increased interferences between the two modalities at 7T (Debener et al., 2008; Jorge et al., 2015b; Mullinger et al., 2008b). We controlled for EEG artifacts by using the artifact acquisition approach of Jorge et al. (2015a), recording 4 electrodes isolated from the scalp to improve data quality. On the other hand, for fMRI, we measured a lower correlation of FC_fMRI_ between the 64Ch-7T dataset and the other datasets (r>0.64 vs. r>0.78 for non-7T datasets, Table 1). This is most likely due to artifacts induced on the fMRI by strong influence of the EEG leads converging to the superior-parietal regions of the cap (Jorge et al., 2015b). This artifact can potentially be avoided by customizing the cabling of the EEG cap (Meyer et al., 2019). We note that this interpretation remains speculative as we did not acquire a proper control for this analysis (fMRI acquisition in the same subjects without EEG-cap in the scanner). Other effects uncontrolled for like TR, scan duration and eyes-closed vs. eyes open might also have a significant effect on the differences between datasets. Future studies should investigate the effects of such improvement on FC_fMRI_.

### Crossmodal relationship and the choice of the spatial resolution of the brain atlas

We observed decreased crossmodal correlation when increasing the number of atlas regions for both the 64-channel EEG setup and the 256-channel setup. Nevertheless FC_EEG-β_ remains the strongest correlation with FCfMRI. This suggests that, when taking an atlas with more and smaller regions, it might be difficult to determine significant differences between frequencies due to lower SNR. Nonetheless, overall, this remains speculative as it also seems that those changes might be driven by the datasets 64Ch1.5T and 256Ch3T, which qualitatively seem to have disproportionally smaller correlations for the Destrieux atlas (SI Table 4). For source reconstruction, the optimal number of distinguishable regions (in terms of cross talk between source reconstructed M/EEG signals) was found to be around 70 regions (Farahibozorg et al., 2018), both for MEG (204 planar gradiometers, 102 magnetometers) and EEG (70 channels). Previous work confirmed a limited improvement in terms of the source reconstruction’s spatial resolution when 256 vs. 64 channels were used (Lantz et al., 2003). Ultimately, our study design cannot formally compare a 64- or 256-channel EEG as the 256-channel cap comes with other setup differences that might also influence the signal(2m cable from amplifier to cap potentially increasing artefacts (Iannotti et al., 2015)). It has been demonstrated that a longer cable length to the amplifier negatively affects the EEG data quality inside the scanner (especially high frequencies such as the gamma band (Jorge et al., 2015b)). Additionally, the electrode/sponge/amplifier system used in the 256-channel setup has generally higher impedances of individual electrodes (e.g. (Foged et al., 2017) used impedance limits of <20kΩ for a 64-electrode electrode/gel system and <50kΩ for the 256-channel electrode/sponge system). Future work should investigate FC_EEG_ as a function of using 64 or 256 electrodes outside the MR-scanner room.

### Low SNR of gamma and the impact of artifacts specific to EEG inside a scanner

Due to the limitation of recording the electrophysiological signal on the scalp, EEG high-γ frequencies (>60Hz) were discarded for analysis. Actually, even the low-γ range from 30-60Hz has been shown to be difficult to analyze in a simultaneous EEG-fMRI setup (Uji et al., 2018). Due to its lower signal power, the EEG-γ band is most affected by a range of different sources of scanner-related artifacts (Jorge et al., 2015b; Uji et al., 2018). As EEG artifacts increase as a function of field strength (Debener et al., 2008), the 64Ch-1.5T performed by far the best in terms of FC_fMRI_-FC_EEG-γ_ correlation (this is despite the drop in BOLD sensitivity, see Table 1). Additionally, the EEG-γ SNR seems to be particularly decreased by long cables (EEG cap to EEG amplifiers) (Jorge et al., 2015b) and vibration artifacts due to the scanner’s helium pump (Jorge et al., 2015b; Nierhaus et al., 2013; Rothlübbers et al., 2015). This is highlighted by our results combining the 256Ch-3T setup with longer cables and the helium pump turned on, which might have caused the additional decrease of FC_EEG-γ_ correlations. An alternative for analyzing the gamma signal at higher field was recently proposed with the use of fast multiband sequences with a fast TR followed by a ‘silent’ period with no scanning to extract the gamma signal (Uji et al., 2018). As another approach to further correct for EEG artifacts in the scanner, the use of electrodes as motion artifact sensors (Jorge et al., 2015a; Masterton et al., 2007) has been proposed to monitor all magnetic induction effects such as gradient, pulse-related (or cardioballistic), vibration and spontaneous motion. Iannotti et al. (2015) proposed a similar approach using cheek electrodes (with less/no neuronal signal) to better estimate the pulse artefact. We demonstrate in our 64Ch-7T data that the motion sensor approach can help to generate topographically reproducible FC_EEG_ even in an artifact-sensitive ultra-high-field scanner setting. Our correction technique used in the 64Ch-7T dataset may have greatly contributed for FC_EEG-γ_ being most correlated to the less artifactual FC_EEG-γ_ recorded in the 1.5T scanner (as compared to the two 3T datasets, Table 2) and FC_fMRI_-FC_EEG-γ_ correlations being higher than in the 64Ch-3T and 256Ch-3T datasets (Fig 2).

### Further methodological considerations and future work

The goal of this study was not to perfectly control for all acquisition and preprocessing steps but to assess the generalizability and reproducibility in a heterogeneous setting, representative of the variability of center specific protocols (Botvinik-Nezer et al., 2020). The reproducible results of the moderate correlation between EEG and fMRI connectivity confirm the feasibility of using this approach in multicentric settings. Further, the measured level of reproducibility of monomodal correlation of connectomes across recording sites determines the potential upper limit of this measure when trying to differentiate (e.g. clinical) outcomes across different datasets (Noble et al., 2019). A measure that does not highly correlate across sites would not be suited for this purpose.

Though brain-wide changes (largest effect size in visual areas) of FC_fMRI_ have been reported between eyes-open and eyes-closed conditions (Agcaoglu et al., 2019). In prior fMRI work, from a test-retest point of view modest increases of reliability are observed for the eyes-open condition (Noble et al., 2019). From a crossmodal point of view Tewarie et al. (2016) reproduced for both eyes-open and eyes-closed condition that MEG and MRI connectomes are correlated (strongest in the beta band consistent with our results). In line with this observation, our crossmodal correlation of EEG and fMRI was comparable in magnitude between eyes-open and eyes closed conditions (SI results). Eyes-open vs. eyes-closed conditions can be interpreted as two separate tasks (Buckner et al., 2013). From this point of view a general stability of ICNs across conditions is also supported by the finding that ICNs are not only observable during rest but also while performing a task (Cole et al., 2014; Krienen et al., 2014).

Future work may study this aspect more specifically by recording both eyes-open and eyes-closed data in the same subjects with one specific EEG-fMRI setup. Equally, alertness and wakefulness have been shown to further confound FC analysis (Tagliazucchi and Laufs, 2014). The described conservative scrubbing of EEG and fMRI and the absence of significant correlation with the crossmodal correlation and the framewise displacement (SI Table 8) speak against head motion as primary contributor to the cross-modal relationship. Nevertheless, there is a possibility that the 72 subjects in our study did not provide sufficient statistical power to measure the potential moderate effect of motion on EEG-fMRI association for delta and gamma bands.

Besides the variability induced by different hardware setups especially for EEG (Pernet et al., 2019), source analysis (Mahjoory et al., 2017) and connectivity estimation (Colclough et al., 2016) provide more heterogeneous analysis options than the more consolidated field of fMRI processing. Better controlling the variable outcome of complex processing pipelines is needed (Botvinik-Nezer et al., 2020; Carp, 2012). Guidelines might recommend specific steps and strategies of best practice to improve neurobiological relevance and reduce erroneous localization/connections (He et al., 2019). Openly available toolkits can further help to streamline this process (Meunier et al., 2020; Schirner et al., 2015).

### Conclusion

In conclusion, we demonstrated the reproducibility of EEG-fMRI connectomes across various acquisition setups and established for the first time the feasibility of extracting EEG-fMRI connectomes at 7T. From an fMRI perspective, the intrinsic connectivity organization of the brain has been linked both to cognitive states and pathology. Reproducible estimation of crossmodal network organization demonstrates the existence of a multimodal functional core and adds a new dimension of how we can assess the healthy and pathologic brain, by dissociating the neurobiological scenarios that may give rise to the observed similarity of FC organization across timescales and modalities.

## Abbreviations

AEC: Amplitude envelope correlation
dMRI: Diffusion Magnet Resonance Imaging
ECG: Electrocardiogram
FC: Functional connectivity
ICA: Independent component analysis
ICN: Intrinsic connectivity network
iCoh: Imaginary part of the coherency
M/EEG: Magneto/Electroencephalography
SC: Structural connectivity

## Acknowledgements

We thank Katia Lehongre, Benjamin Morillon (64Ch-3T dataset) and Fani Deligianni and Jonathan Clayden (64Ch-1.5T dataset) for generously sharing their data. We acknowledge support from Swiss National Science Foundation (SNSF, under grants CRSII5_170873, 169198 and 192749 to SV and 188769 to FG). SS acknowledges financial support from NIH R01MH11622601A1 and R21NS10460302. ALG was supported by ERC 260347 – COMPUSLANG. JJ and RG were supported by Centre d’Imagerie BioMédicale (CIBM) of the UNIL, UNIGE, HUG, CHUV, EPFL and the Leenaards and Jeantet Foundations.

## Data Availability

All connectomes are available on Zenodo (https://doi.org/10.5281/zenodo.3905103). The 64Ch-1.5T raw data is publicly available at https://osf.io/94c5t/. The other raw data will be made available by request to SV (256Ch-3T), ALG (64Ch-3T) and JJ (64Ch-7T).

## Supplementary Information

**SI Table 1:** Summary of selected acquisition parameters and data analysis steps

|  | 64Ch-1.5T | 64Ch-3T | 256Ch-3T | 64Ch-7T |
| --- | --- | --- | --- | --- |
| **Subjects** |  |  |  |  |
| Number | 16 | 26 | 21 | 9 |
| Mean Age | 32.87 | 24.39 | 32.13 | 23.56 |
| Females | 6 | 8 | 8 | 4 |
| **Paradigm** |  |  |  |  |
| Eyes-open/closed | Open | Closed | Closed | Open |
| Duration | 10min48s | 30 min | 4min58.5s | 8min |
| **Hardware stack** |  |  |  |  |
| Scanner | Siemens Avanto | Siemens Tim-Trio | Siemens Magnetom Trio | Siemens Magnetom |
| TR | 2160ms | 2000ms | 1990ms | 1000ms |
| Helium Pump | Off | Off | On | On |
| Electrodes | 64 (63 active) | 64 (62 active) | 256 (204 active) | 64 (59 active) |
| Amplifier | Behind the scanner (next to the head of the participant) | Behind the scanner | In front of the scanner (next to the feet of the participant) | Behind the scanner |
| Impedance | <100kΩ | <20kΩ | <50 kΩ | <20kΩ |
| **Software stack** |  |  |  |  |
| Gradient artifact | BrainVisionAnalyser 2 | FMRIB | Inhouse (Iannotti et al., 2015) | Inhouse (Jorge et al., 2015a) |
| Pulse artifact | BrainVisionAnalyser 2 | FMRIB | Inhouse (Iannotti et al., 2015) | Inhouse (Jorge et al., 2015a) |
| ICA cleaning | Yes (BrainVisionAnalyser 2) | No | Yes (EEGLab, runICA v1.29) | Yes (inhouse ICA ext. Infomax algorithm) |
| Brainstorm | 10<sup>th</sup> August 2017 | 10<sup>th</sup> August 2017 | 15<sup>th</sup> January 2019 | 15<sup>th</sup> January 2019 |
| Freesurfer | v6.0.0 | v6.0.0 | v6.0.0 | v6.0.0 |
| SPM | SPM12 (revision 6906) | SPM12 (revision 6906) | SPM12 (revision 7475) | SPM12 (revision 7475) |
| FSL | v5.0 | v5.0 | v5.0 | V6.0 |
| ANTs | N/A | N/A | N/A | V2.2.0 |

**SI Table 2:**
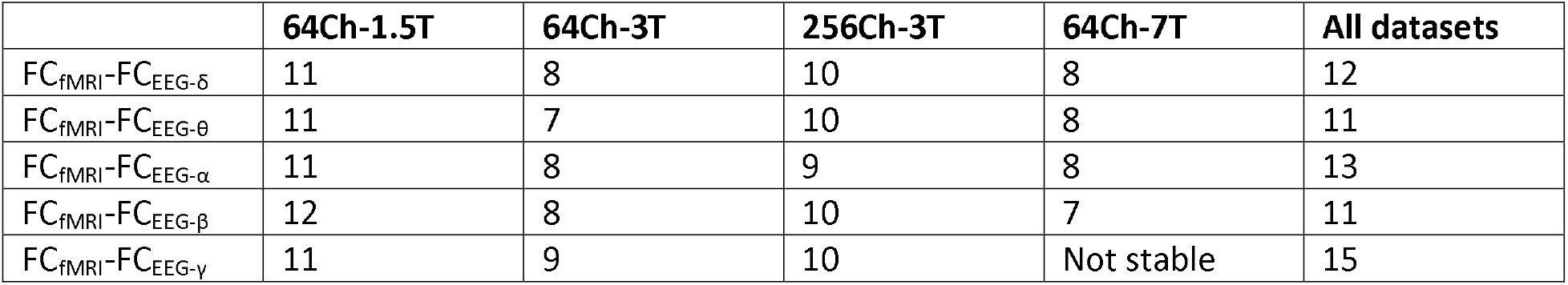
The number of subjects needed to generate a stable crossmodal correlation estimate. The stability is assessed by adding a random subject to the average and evaluating the change in crossmodal correlation. The crossmodal correlation is considered stable when the change is not more than 1% of the crossmodal correlation of this frequency band derived from all 72 subjects. Note that for the 64Ch-7T dataset for FC_EEG-γ_ no stability was reached when talking all of the 9 subjects.

**SI Table 3:** P-values of permutation tests comparing the dataset averaged crossmodal FC_fMRI_-FC_EEG_ correlation between different EEG frequency bands (5000 iterations/512 iterations for 64Ch-7T dataset, Bonferroni-corrected significance level p<0.05 which corresponds to the uncorrected level p<0.05/50 = 0.001, significant cells are marked in bold). Note that the 95% percentile binominal proportion confidence interval of the permutation test is given by: p ± 1.96√(p(1-p)/n) with p being the estimated p-value and n being the number of iterations. For a p-value at Bonferroni-threshold the interval is 0.001 ± 0.00088 (5000 iterations) and 0.001 ± 0.00274 (512 iterations).

|  | 64Ch-1.5T | 64Ch-3T | 256Ch-3T | 64Ch-7T | All datasets |
| --- | --- | --- | --- | --- | --- |
| $\beta > \delta$ | **0.0002** | 0.0102 | **0.0002** | 0.0078 | **<0.0002** |
| $\beta > \theta$ | 0.0014 | 0.0186 | **<0.0002** | 0.1641 | **<0.0002** |
| $\beta > \alpha$ | 0.0036 | 0.0044 | 0.1918 | 0.1335 | **<0.0002** |
| $\beta > \gamma$ | 0.1254 | **<0.0002** | **<0.0002** | 0.0215 | **<0.0002** |
| $\alpha > \delta$ | 0.2672 | 0.6978 | 0.0022 | 0.1563 | 0.0172 |
| $\alpha > \theta$ | 0.166 | 0.5862 | 0.0198 | 0.6172 | 0.075 |
| $\alpha > \gamma$ | 0.8546 | **0.0002** | **0.0004** | 0.6016 | 0.0084 |
| $\theta > \delta$ | 0.6596 | 0.6948 | 0.0044 | 0.1797 | 0.0816 |
| $\theta > \gamma$ | 0.9662 | 0.0094 | 0.0026 | 0.3859 | 0.0752 |
| $\delta > \gamma$ | 0.951 | 0.0012 | 0.023 | 0.8926 | 0.201 |

**SI Table 4:**
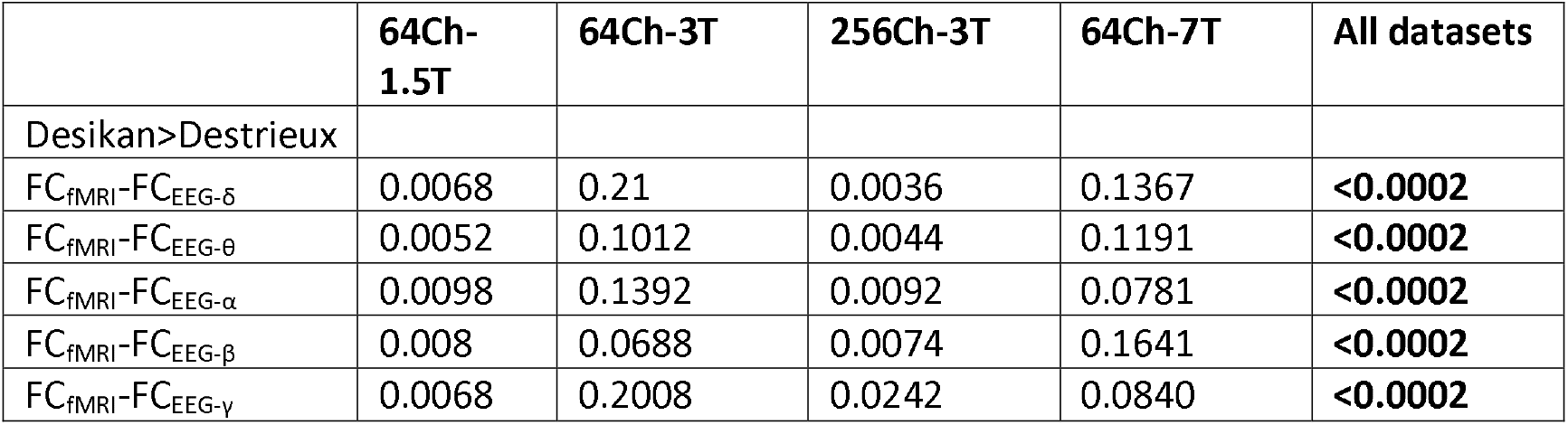
P-values of permutation tests comparing the dataset averaged crossmodal FC_fMRI_-FC_EEG_ correlation between different atlases (5000 iterations/512 iterations for 64Ch-7T dataset, Bonferroni-corrected significance level p<0.05 which corresponds to the uncorrected level p<0.05/25 = 0.002, significant cells are marked in bold). Note that the 95% percentile binominal proportion confidence interval of the permutation test is given by: p ± 1.96√(p(1-p)/n), with p being the estimated p-value and n being the number of iterations. For a p-value at Bonferroni-threshold the interval is 0.002 ± 0.00124 (5000 iterations) and 0.002 ± 0.00387 (512 iterations).

|  | 64Ch-1.5T | 64Ch-3T | 256Ch-3T | 64Ch-7T | All datasets |
| --- | --- | --- | --- | --- | --- |
| Desikan>Destrieux |  |  |  |  |  |
| $FC_{fMRI}-FC_{EEG-\delta}$ | 0.0068 | 0.21 | 0.0036 | 0.1367 | **<0.0002** |
| $FC_{fMRI}-FC_{EEG-\theta}$ | 0.0052 | 0.1012 | 0.0044 | 0.1191 | **<0.0002** |
| $FC_{fMRI}-FC_{EEG-\alpha}$ | 0.0098 | 0.1392 | 0.0092 | 0.0781 | **<0.0002** |
| $FC_{fMRI}-FC_{EEG-\beta}$ | 0.008 | 0.0688 | 0.0074 | 0.1641 | **<0.0002** |
| $FC_{fMRI}-FC_{EEG-\gamma}$ | 0.0068 | 0.2008 | 0.0242 | 0.0840 | **<0.0002** |

**SI Table 5:**
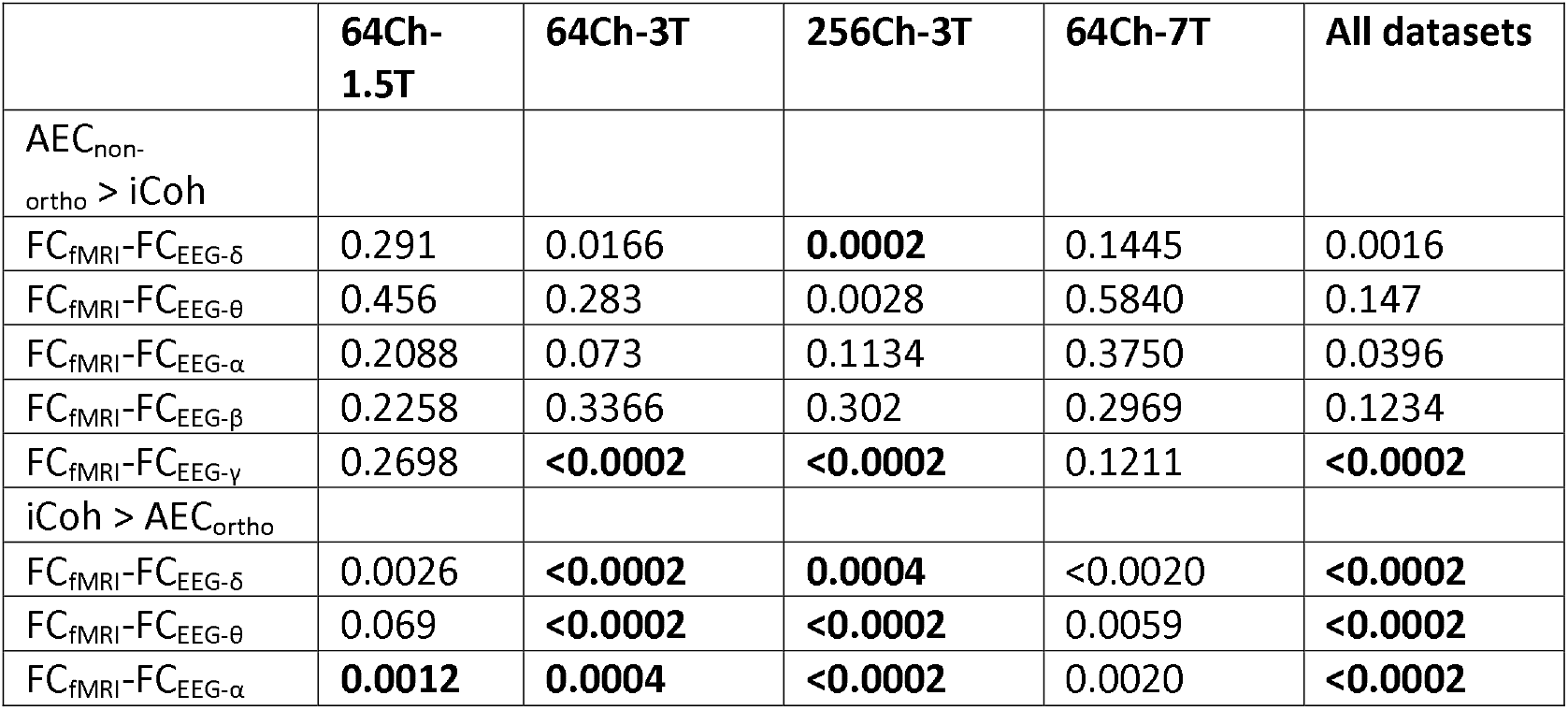
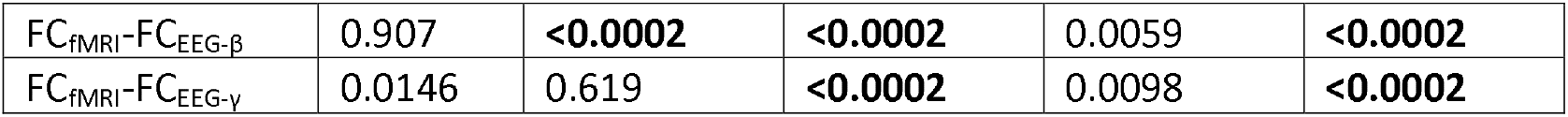
P-values of permutation tests comparing the dataset averaged crossmodal FC_fMRI_-FC_EEG_ correlation between different EEG connectivity measures (5000 iterations/512 iterations for 64Ch-7T dataset, Bonferroni-corrected significance level p<0.05 which corresponds to the uncorrected level p<0.05/50 = 0.001, significant cells are marked in bold). Note that the 95% percentile binominal proportion confidence interval of the permutation test is given by: p ± 1.96√(p(1-p)/n), with p being the estimated p-value and n being the number of iterations. For a p-value at Bonferroni-threshold the interval is 0.001 ± 0.00088 (5000 iterations) and 0.001 ± 0.00274 (512 iterations).

**SI Table 6:** P-values of permutation tests comparing the dataset averaged crossmodal FC_fMRI_-FC_EEG_ correlation using the complete subject session vs. the first 4min58.5s and using the average connectivity over all subjects vs. the first 9 subjects (5000 iterations/512 iterations for 64Ch-7T dataset, Bonferroni-corrected significance level p<0.05 which corresponds to the uncorrected level p<0.05/20 = 0.0025, significant cells are marked in bold). Note that the 95% percentile binominal proportion confidence interval of the permutation test is given by: p ± 1.96√(p(1-p)/n), with p being the estimated p-value and n being the number of iterations. For a p-value at Bonferroni-threshold the interval is 0.0025 ± 0.00138 (5000 iterations) and 0.0025 ± 0.00433 (512 iterations).

|  | 64Ch-1.5T | 64Ch-3T | 256Ch-3T | 64Ch-7T | All datasets |
| --- | --- | --- | --- | --- | --- |
| All session > truncated to 4min58.5 |  |  |  |  |  |
| $FC_{fMRI}-FC_{EEG-\delta}$ | 0.0472 | 0.1398 | | 0.7715 | 0.1418 |
| $FC_{fMRI}-FC_{EEG-\theta}$ | 0.041 | 0.2178 | | 0.6621 | 0.132 |
| $FC_{fMRI}-FC_{EEG-\alpha}$ | 0.1594 | 0.2748 | | 0.4570 | 0.2088 |
| $FC_{fMRI}-FC_{EEG-\beta}$ | 0.0758 | 0.1558 | | 0.8906 | 0.2146 |
| $FC_{fMRI}-FC_{EEG-\gamma}$ | 0.0312 | 0.0794 | | 0.8555 | 0.0724 |
| All subjects > 9 subjects |  |  |  |  |  |
| $FC_{fMRI}-FC_{EEG-\delta}$ | 0.0682 | 0.192 | 0.0678 | | 0.0116 |
| $FC_{fMRI}-FC_{EEG-\theta}$ | 0.015 | 0.2226 | 0.0814 | | 0.0076 |
| $FC_{fMRI}-FC_{EEG-\alpha}$ | 0.0388 | 0.254 | 0.0292 | | 0.0056 |
| $FC_{fMRI}-FC_{EEG-\beta}$ | 0.0064 | 0.2548 | 0.0682 | | 0.0098 |
| $FC_{fMRI}-FC_{EEG-\gamma}$ | **<0.0002** | 0.0938 | 0.0212 | | **0.0006** |

**SI Table 7:** P-values of t-test comparing the crossmodal correlation of each subjects between derived from the entire session and from the first 4min58.5 (one-sided ttest, Bonferroni-corrected significance level p<0.05 which corresponds to the uncorrected level p<0.05/15 = 0.0033, significant cells are marked in bold). Note that the 95% percentile binominal proportion confidence interval of the permutation test is given by: p ± 1.96√(p(1-p)/n), with p being the estimated p-value and n being the number of iterations. For a p-value at Bonferroni-threshold the interval is 0.0033 ± 0.00159 (5000 iterations) and 0.0033 ± 0.00497 (512 iterations).

|  | 64Ch-1.5T | 64Ch-3T | 64Ch-7T |
| --- | --- | --- | --- |
| All session > truncated to 4min58.5 |  |  |  |
| $FC_{fMRI}-FC_{EEG-\delta}$ | 0.1196 | **$6.4 \cdot 10^{-06}$** | 0.5291 |
| $FC_{fMRI}-FC_{EEG-\theta}$ | 0.1597 | **0.0003** | 0.6987 |
| $FC_{fMRI}-FC_{EEG-\alpha}$ | 0.3196 | 0.0199 | 0.7764 |
| $FC_{fMRI}-FC_{EEG-\beta}$ | 0.0592 | **0.0002** | 0.9533 |
| $FC_{fMRI}-FC_{EEG-\gamma}$ | 0.1023 | 0.0352 | 0.9631 |

### Impact of movement

**SI Table 8:** To assess the impact of movement on the individual connectome, we correlated the individual crossmodal correlation with the mean framewise displacement derived from fMRI. We did not find any significant correlations between the two measures, although qualitatively moderate-sized associations were observed between FD and the EEG-fMRI crossmodal correlation for delta and gamma bands (Bonferroni-corrected for 5 frequencies, significance level p<0.05 corresponds to p<0.05/5=0.01).

|  | r | p |
| --- | --- | --- |
| $FC_{fMRI}-FC_{EEG-\delta}$ vs. FD | -0.2440 | 0.0389 |
| $FC_{fMRI}-FC_{EEG-\theta}$ vs. FD | -0.1674 | 0.1599 |
| $FC_{fMRI}-FC_{EEG-\alpha}$ vs. FD | -0.0492 | 0.6812 |
| $FC_{fMRI}-FC_{EEG-\beta}$ vs. FD | -0.1607 | 0.1774 |
| $FC_{fMRI}-FC_{EEG-\gamma}$ vs. FD | -0.2944 | 0.0121 |

### Impact of eyes-open vs. eyes-resting state

It has been shown by Mo et al. (2013) that EEG alpha power is coupled to DMN activity when alternating between an eyes-open and an eyes-closed paradigm. We tested if we could observe any systematic changes of the crossmodal correlation between eyes-open and eyes-closed condition by permuting the labels of the different datasets (5000 permutations). As we did not measure eyes-open condition in the same subjects and setup these results should not be overinterpreted. We did not observe any significant changes of crossmodal correlation between both conditions in any frequency band (p>0.05) except for the FC_fMRI_-FC_EEG-γ_ correlation (p<0.0002). Those changes in FC_fMRI_-FC_EEG-γ_ most likely stem from the general higher crossmodal correlation of FC_fMRI_-FC_EEG-γ_ for the datasets of the eyes-open condition (64Ch-1.5T and 64Ch-7T, see Fig 2). In order to exclude that the spatial contributions to the crossmodal correlation in the visual network were only driven by eyes-open ore eyes closed effects we recalculated the spatial contribution for each dataset. We observe that the significant spatial contribution in the visual network is present in all frequencies and in all datasets but the FC_fMRI_-FC_EEG-α_ of the 64Ch-3T dataset (SI Table 9). As such we did not observe any systematic connectivity differences between eyes-open datasets (64Ch-1.5T and 64Ch-7T) and eyes closed datasets (64Ch-3T and 256Ch-3T), especially for FC_fMRI_-FC_EEG-α_. This is in line with our previous results when analyzing FC dynamics (Wirsich et al., 2020b). We conclude that the FC_fMRI_-FC_EEG_ crossmodal correlation might be primarily capturing the intrinsic coupling networks (ICN) linked to canonical ICNs (Yeo et al., 2011), which are preserved across eyes-open vs. eyes closed resting state.

**SI Table 9:** P-values when comparing the spatial contribution of the visual network connections to the crossmodal FC_fMRI_-FC_EEG_ correlation as compared to inter-ICN connections for each frequency band and dataset (one-sided t-test spatial contribution Visual>interICN: Bonferroni correction threshold for 5 frequencies and 4 datasets is defined at p=0.05/20=0.0025).

| VIS>interICN | 64Ch-1.5T | 64Ch-3T | 256Ch-3T | 64Ch-7T |
| --- | --- | --- | --- | --- |
| $FC_{fMRI}-FC_{EEG-\delta}$ | $1.5 \cdot 10^{-23}$ | $1.1 \cdot 10^{-11}$ | $2.1 \cdot 10^{-83}$ | $2.4 \cdot 10^{-37}$ |
| $FC_{fMRI}-FC_{EEG-\theta}$ | $1.6 \cdot 10^{-21}$ | $1.4 \cdot 10^{-6}$ | $4.5 \cdot 10^{-62}$ | $2.6 \cdot 10^{-50}$ |
| $FC_{fMRI}-FC_{EEG-\alpha}$ | $1.2 \cdot 10^{-120}$ | 0.0376 | $4.2 \cdot 10^{-89}$ | $5.1 \cdot 10^{-85}$ |
| $FC_{fMRI}-FC_{EEG-\beta}$ | $3.9 \cdot 10^{-63}$ | $3.2 \cdot 10^{-21}$ | $2.3 \cdot 10^{-54}$ | $4.5 \cdot 10^{-62}$ |
| $FC_{fMRI}-FC_{EEG-\gamma}$ | $2.6 \cdot 10^{-69}$ | $7.6 \cdot 10^{-87}$ | $1.7 \cdot 10^{-58}$ | $1.2 \cdot 10^{-26}$ |

